# Introduction of a condensed, reverse tricarboxylic acid cycle for additional CO_2_ fixation in plants

**DOI:** 10.1101/2022.03.04.483018

**Authors:** Nathan J. Wilson, Caroline M. Smith-Moore, Yuan Xu, Brianne Edwards, Christophe La Hovary, Swathi Barampuram, Kai Li, Denise Aslett, Mikyoung Ji, Xuli Lin, Simina Vintila, Manuel Kleiner, Deyu Xie, Yair Shachar-Hill, Amy Grunden, Heike Sederoff

## Abstract

Plants employ the Calvin-Benson cycle (CBC) to fix atmospheric CO_2_ for the production of biomass. The flux of carbon through the CBC is limited by the activity and selectivity of ribulose-1,5-bisphosphate carboxylase/oxygenase (RuBisCO). Alternative pathways that do not use RuBisCO to fix CO_2_ exist but occur only in anaerobic microorganisms. Rather than modifying existing routes of carbon metabolism in plants, we have developed a synthetic carbon fixation cycle that does not exist in nature, but is inspired by metabolisms of bacterial autotrophs. This synthetic cycle uses endogenous plant metabolites to fix CO_2_ and yield glyoxylate as a product. In this work, we build and characterize a condensed, reverse tricarboxylic acid (crTCA) cycle *in vitro* and *in planta*. We demonstrate that a simple, synthetic cycle can be used to fix carbon *in vitro* under aerobic and mesophilic conditions and that these enzymes retain activity when expressed transiently *in planta*. We then evaluate stable transgenic lines of *Camelina sativa* that have both phenotypic and physiologic changes. Transgenic *C. sativa* are shorter than controls with increased rates of photosynthetic CO_2_ assimilation and changes in photorespiratory metabolism. This first iteration of a build-test-learn phase of the crTCA cycle provides promising evidence that this pathway can be used to increase photosynthetic capacity in plants.

## Introduction

Within the past decade, there has been great interest in increasing the photosynthetic efficiency in plants. This is due to the growing need for increased yields in agriculture as the world population continues to increase, a changing climate results in extreme weather events and agricultural yields stagnate [1, 2]. The photosynthetic process is a complex series of chemical reactions that derive their energy from sunlight to fix and store carbon from the atmosphere. Improvements to this process are regarded as an achievable route to increasing overall agricultural yields [2–4]. In C3 plants, the enzyme RuBisCO is the entry point of CO_2_ into plant metabolism. This enzyme, however, is “notoriously inefficient” due to its low catalytic rate (∼1– 10 CO_2_ molecules s^-1^) and its oxygenase activity [5]. The oxygenase reaction of RuBisCO necessitates the energetically costly process of photorespiration, by which the plants lose up to an estimated 30% of the energy gained from photosynthesis. Under warmer climate conditions, RuBisCO is more likely to participate in oxygenase activity making improvement of this inefficiency an imperative under future climate predictions [6].

Attempts at engineering RuBisCO directly are hampered by the multifaceted layers of regulation, biogenesis/assembly factors and kinetic considerations associated with the enzyme [7–9]. An emerging field of research is the use of synthetic metabolisms to further redirect or enhance plant metabolism [10]. Several synthetic approaches to decrease or bypass photorespiratory metabolism have been successful in several species of plants including *Arabidopsis* [11–13], tobacco [14, 15], *Camelina* [16] and rice [17]. Alternatively, an emerging field of interest lies in designing metabolisms that are capable of supplementing natural CO_2_ fixation. Synthetic carbon fixation pathways are theoretical means to circumvent or “bypass” the inefficiencies associated with RuBisCO [18]. Non-photosynthetic autotrophic bacteria have evolved RuBisCO-independent CO_2_ fixation pathways and may offer ways to enhance CO_2_ fixation in plants [18, 19]. The first example of a synthetic carbon fixation pathway to be realized *in vitro* was the CETCH cycle. This cycle used 13 core enzymes, several of which received active site optimization, along with 4 additional co-enzymes to power the cycle. These new-to-nature metabolisms have increased carboxylation capacities and use less cellular energy than existing pathways and represent novel ways to enhance plant physiology [20].

Bar-Even, et al. [18] computationally evaluated several theoretical synthetic cycles that fix CO_2_. The shortest theoretical carbon fixation pathway posited by these authors was a modification of the reverse TCA cycle, which could be modified to be energetically feasible under diverse physiological conditions. In this study, we construct a condensed, reverse/reductive TCA cycle (crTCA) cycle based on the original design first postulated by Bar-Even et al. [18]. Five enzymes make up the core of the crTCA cycle: 1) succinyl CoA synthetase (SCS, E.C. 6.2.1.5) which converts succinate into succinyl CoA with the use of one ATP; 2) 2-oxoglutarate:ferredoxin oxidoreductase (KOR, E.C. 1.2.7.3) carboxylates the succinyl CoA into 2-oxoglutarate by oxidizing ferredoxin; 3) 2-oxoglutarate carboxylase (OGC, E.C. 6.4.1.7) subsequently carboxylates the 2-oxoglutarate to produce oxalosuccinate at the cost of one ATP; 4) oxalosuccinate reductase (OSR, sometimes referred to as isocitrate dehydrogenase or ICDH, E.C. 1.1.1.42) uses one NADPH to reduce oxalosuccinate to isocitrate; and 5) isocitrate lyase (ICL, E.C. 4.1.3.) hydrolyzes the isocitrate to glyoxylate (the cycle’s product) and succinate which serves as substrate for another iteration of the cycle (**Fig 1**). The initial design and analysis of such a cycle indicated that while a modified reverse TCA cycle is the most simple (fewest enzymatic steps), the thermodynamics of several reactions are not favorable at physiologic conditions/concentrations. This situation is further complicated because the use of a modified reverse TCA cycle depends on a class of enzymes known as 2-oxoglutarate:ferredoxin oxidoreductases (KORs) – which are often oxygen sensitive [18]. In this work, however, we have identified enzymes that are capable of functioning as a metabolic cycle *in vitro* at aerobic, mesophilic conditions. We then installed the crTCA cycle in the chloroplasts of the biofuel crop *Camelina sativa* to assess if the crTCA cycle can functionally enhance net CO_2_ assimilation and increase plant productivity.

**Fig 1:**
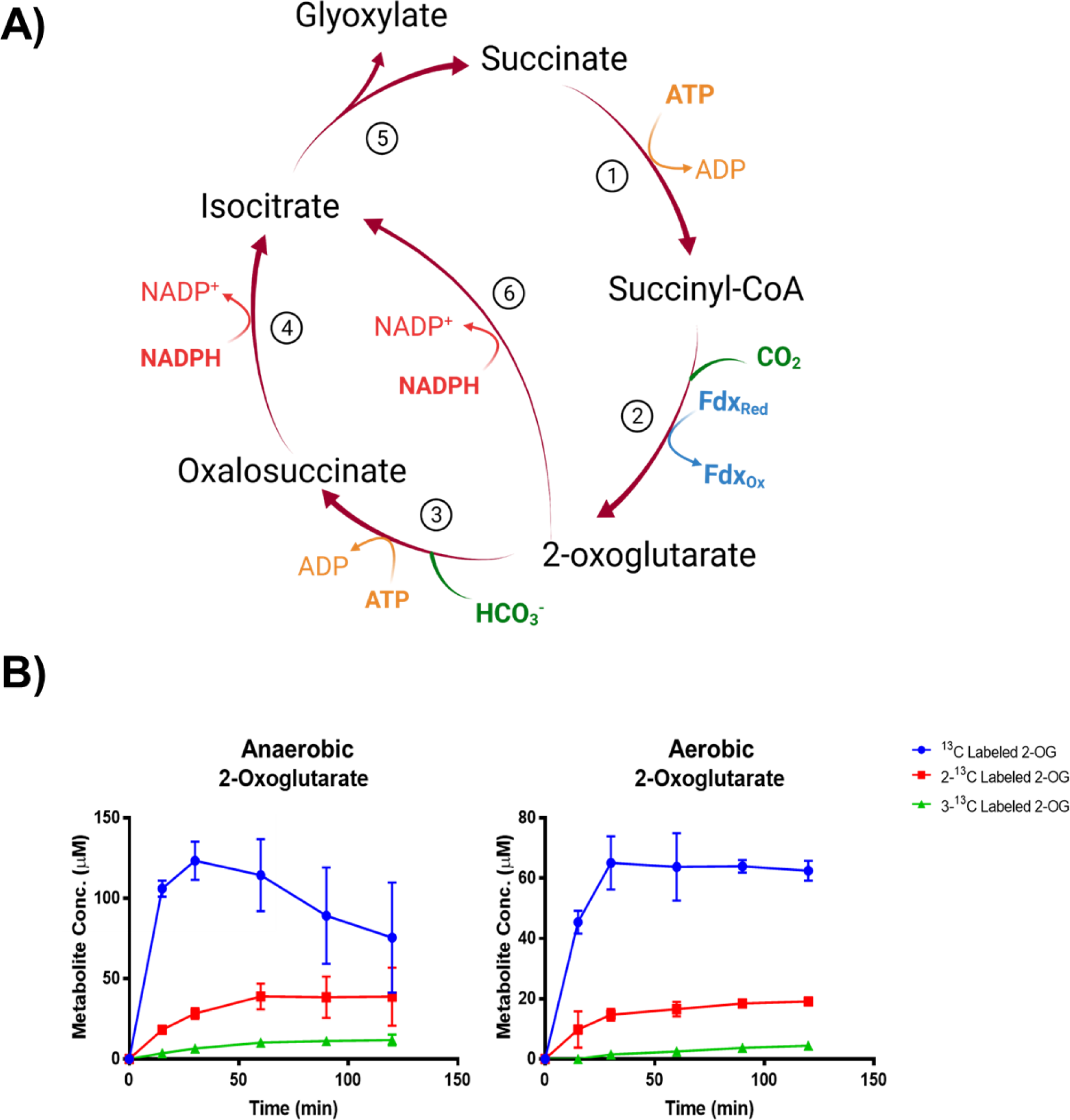
The crTCA cycle, selected enzymes and *in vitro activity.* A) Schematic representation of a condensed, reverse TCA (crTCA) cycle. The design of the cycle is to take succinate through four or five steps to yield glyoxylate and fixing two carbon molecules. Reactions listed by number are 1) succinyl-CoA synthetase (SCS); 2) 2-oxoglutarate:ferredoxin oxidoreductase (KOR); 3) 2-oxoglutarate carboxylase (OGC); 4) oxalosuccinate reductase (OSR); 5) isocitrate lyase (ICL); 6) isocitrate dehydrogenase (ICDH). B) Enzyme reactions were provided with NaH^13^CO_3_ as substrate and samples were taken over the course of two hours. Isotope incorporation into the metabolites were quantified using standard curves analyzed at the same time as the reaction samples. The mean is shown ± one standard deviation, n = 3.

## Results

### Selection of enzymes used in the crTCA cycle

A condensed, reversed TCA cycle can be conceptually achieved through four or five enzymatic steps [18]. In this work, the selection of five candidate enzymes was based on sequence alignment to functionally characterized enzymes, activity predictions from sequence, and the physiology of the source bacteria (e.g. mesophilic, aerobic). Four to seven enzyme candidates were selected for each of the crTCA cycle steps (**Supplementary Table S1**). Three out of 4 SCSs, 4 out of 6 KORs, 4 out of 7 OGCs, 3 out of 5 OSRs, and 4 out of 5 ICLs were purified with a yield of more than 5 mg/L of *E. coli* culture.

The purified proteins were first tested for activity using UV-Vis spectroscopy methods (**Table 1**). For SCS, OGC, and ICL, UV-Vis spectrophotometric methods measured the reaction rate in the forward direction. SCS from *Bradyrhizobium sp.* BTAi1 and ICL from *Nocardia farcinica* IFM 10152 were selected for their specific activities. While OGC from *Mariprofundus ferrooxydans* PV-1 had a similar specific activity to other overexpressed OGC enzymes, its protein yield was much higher. For OSR, UV-Vis spectrophotometry was used to measure the reaction in the reverse direction of the crTCA cycle, due to the lability of the reactant metabolite, oxalosuccinate. Therefore, the recombinant OSRs were initially screened using UV-Vis spectrophotometry and then LC-MS was used to determine the best combination of OGC and OSR. OSR from *Nitrosococcus halophilus* Nc4 coupled with the OGC from *M. ferrooxydans* PV-1 was the best choice for our crTCA cycle under experimental conditions. OSR enzymes are often capable of catalyzing 2-OG carboxylation, performing both the carboxylation of 2-oxoglutarate to oxalosuccinate and reducing the oxalosuccinate to isocitrate. These enzymes are known as isocitrate dehydrogenases (ICDH) [21–24]. As such, NiHa OSR was evaluated for carboxylation activity using a UV-Vis assay measuring the oxidation of NAD^+^ [25]. This OSR enzyme was found to carboxylate 2-OG and therefore serves as an ICDH (**Supplementary Table S2**).

**Table 1:** The studied crTCA enzymes and *in vitro*, *in planta* activities. The enzymes used in this study for the realization of a crTCA cycle. Size determination for each subunit (α or β) is estimated from sequence data. Activity for each enzyme was determined using a combination of UV-Vis and LC-MS based activity assays. The mean is shown ± one standard deviation, n = 3. The NiHa OSR (Step 4) enzyme was shown to have activity as both an oxalosuccinate reductase and an isocitrate dehydrogenase. *In planta* activity shown is the mean of three biological replicates ± one standard deviation.

|  | Source Organism | Size (kDa) | *In vitro* activity (nmol min<sup>-1</sup> mg<sup>-1</sup>) | *In planta* activity (nmol min<sup>-1</sup> mg<sup>-1</sup>) |
| --- | --- | --- | --- | --- |
| 1: Succinyl CoA synthetase (SCS) | *Bradyrhizobium* sp.BTAi1 (BrBt) | $\alpha$: 43<br>$\beta$: 30 | 23.7 ± 0.5 | 29.5 + 5 |
| 2: 2-oxoglutarate:ferredoxin oxidoreductase (KOR) | *Hydrogenobacter thermophilus* TK-6 (HyTh) | $\alpha$: 65<br>$\beta$: 31 | 0.22 ± 0.03 | 0.14 + 0.06 |
| 3: 2-oxoglutarate carboxylase (OGC) | *Mariprofundus ferrooxydans* PV-1 (MaFe) | $\alpha$: 53<br>$\beta$: 68 | 0.032 ± 0.002 | N.D. |
| 4: Oxalosuccinate reductase (OSR) | *Nitrosococcus halophilus* Nc4 (NiHa) | 47 | 19 ± 1 (oxalosuccinate)<br>0.24 ± 0.01 (2-oxoglutarate) | 5.8 + 0.44 (oxalosuccinate) |
| 5: Isocitrate lyase (ICL) | *Nocardia farcinica* IFM 10152 (NoFa) | 48 | 10.0 ± 0.3 | 2.42 + 0.25 |

Experiments using KOR were initially performed under anaerobic conditions unless otherwise specified. The KOR enzyme requires reduced ferredoxin for the carbon fixation reaction, which converts the four-carbon derivative, succinyl-CoA to the five-carbon product, 2-OG. The KOR from *Bacillus sp.* M3-13 (BaM3 KOR) showed the highest specific activity for the reverse reaction at RT, however attempts to run the full cycle with this enzyme did not succeed. Instead a combination of KOR and ferredoxin from *Hydrogenobacter thermophilus* TK-6 [26] showed a greater preference for the forward reaction. Despite initial concerns about activity at mesophilic temperatures, the HyTh KOR was evaluated for activity at lower temperatures and was found to have activity at ambient temperature. Thus, the KOR and ferredoxin (FDX) from *H. thermophilus* TK-6 were selected. The best candidates for each crTCA step are listed in **Table 1**.

### *In vitro* demonstration of cyclic carbon fixation by the crTCA cycle

After confirming individual activity, cyclic carbon fixation of the crTCA enzymes was first shown *in vitro* when performed under anaerobic conditions. In this assay, 2-OG and succinyl-CoA were used as the starting metabolites and NaH^13^CO_3_ was used as the carbon source. Multiple ^13^C-labeled glyoxylic acid, succinic acid and 2-OG were detected (**Supplementary Table S3**), establishing that the crTCA cycle fixes carbon under tested conditions. As expected, the amount of multiple labeled metabolites increases with longer reaction times. The succinic acid detected by LC-MS in this experiment is the total amount of free succinic acid plus succinyl-CoA because the LC-MS method was unable to distinguish them.

As KOR enzymes are oxygen sensitive [18], it is important to demonstrate that the crTCA cycle can function under aerobic conditions, such as those present in the plant chloroplast. To test the full function of the crTCA cycle under aerobic conditions, the previous experiment was replicated under aerobic conditions. The production of labeled 2-OG was estimated because KOR is responsible for the carboxylation of succinyl-CoA to produce 2-OG. From LC-MS, single, double, and triple ^13^C-labeled 2-OG were detected (**Fig 1B**). The amount of these metabolites is less than that produced in the anaerobic samples; however, their detection confirms that the crTCA cycle can function in the presence of oxygen. The maximum amounts of ^13^C-labeled metabolites detected under both atmospheric conditions are summarized in **Supplementary Table S3**.

Most of the crTCA cycle enzymes are theoretically capable of catalyzing the reverse reaction to their proposed cycle activity [18]. To evaluate this potential, ICL and OSR were combined to create a coupled reaction. This reaction evaluated the ability of ICL to utilize glyoxylate and succinate to produce isocitrate. The resulting isocitrate would be decarboxylated by OSR to produce 2-oxoglutarate. The specific activity of this reaction was 0.07 ± 0.01 U mg^-1^ based on three replicates. While the measured activity is low compared to the activity of ICL in the production of glyoxylate and succinate (**Table 1**), the ICL and OSR can function together in reverse to produce 2-OG. To test whether the full cycle can also proceed in the reverse direction, succinate and glyoxylate were used as starting reagents with all of the crTCA cycle enzymes. The amount of glyoxylate decreases significantly after incubating 60 minutes at 30° C, while little change was found for the control samples with no succinate or no ICL (**Supplementary Fig S1**). On the contrary, a large amount of 2-OG was detected in the full cycle reactions, suggesting 2-OG was produced under the experimental conditions. This consumption of glyoxylate and production of 2-OG indicate that the crTCA cycle can proceed *in vitro* in the decarboxylation direction at room temperature and aerobic conditions.

To verify the activity of the OSR/ICDH enzyme from *Nitrosococcus halophilus,* we compared the 4-step (SCS, KOR, OSR and ICL) vs. the 5-step (SCS, KOR, OSR/ICDH, OGC and ICL) crTCA cycle. In the four-step cycle OGC was eliminated and OSR would be relied on to perform the carboxylation of 2-OG to produce isocitrate. The reactions were performed and analyzed by LC-MS. The peak areas were compared for the unlabeled and labeled metabolites and showed no statistically significant difference between the two reactions (**Supplementary Fig S2**), suggesting the crTCA cycle may actually perform primarily as a four-step cycle.

### In planta expression of the crTCA cycle

The target application of our synthetic cycle is to improve the carbon fixation capacity of C3 plants by expressing the selected crTCA cycle enzymes in the chloroplast. To test if plants can produce active crTCA cycle enzymes, we transiently transformed either *N. tabacum* or *N. benthamiana* leaves with the individual crTCA cycle genes [27]. The expression of the crTCA cycle enzymes was detected using western blots (**Supplementary Fig S3**). Detection of KOR required expression in *N. benthamiana* instead of *N. tabacum,* as no KOR was detected using *N. tabacum*. To confirm functional expression, enzyme activity was measured in protein extracted from the tobacco leaf tissue. *In planta* activity was found for the four core enzymes with OGC having undetectable carboxylation activity (**Table 1**).

The design of the crTCA cycle consists of five enzymes. Three of these enzymes (SCS, KOR and OGC) are multi-subunit and KOR and OGC require co-enzymes (a ferredoxin and biotin lyase, respectively). We cloned the ten ORFs into three separate vectors (**Supplementary Fig S4, Supplementary Table S4**). The expression of each coding sequence of the crTCA genes is controlled by a constitutive promoter and is also preceded by a chloroplast targeting peptide (ctp). Plants confirmed via PCR for the presence of all three transgenic vectors in their genomes are considered full crTCA lines. These lines were taken to homozygosity through various generations of segregation and analysis. Each generation of the crTCA lines in *C. sativa* were evaluated for transgenic expression using either qRT-PCR or RNA sequencing. The expression of individual transgenes varied greatly across individuals even when the same promoter was used. Most of the crTCA transcripts were present at relatively the same or slightly higher abundances compared to the endogenous ubiquitin (UBQ) control (**Supplementary Fig S5**). It was found, however, that the transcript abundance of the nuclear RbcS, the small subunit of RuBisCO, was approximately 100-200 times greater (**Supplementary Fig S5**). We then used proteomics to measure relative abundances of crTCA proteins in leaf tissue and isolated chloroplasts. In the transgenic lines expressing all five enzymes, we only detected three of the five crTCA proteins (OSR/ICDH, ICL and SCS). In a partial cycle line (“Construct 1”) expressing only three enzymes, we detected all three proteins (OSR/ICDH, ICL and OGC), however the LSU of the OGC enzyme was below the threshold of detection. In comparison, RbcS was approximately 500-1000 times more abundant (**Supplementary Table S5**).

### Transgenic *C. sativa* exhibit changes to morphology and photosynthesis

Transgenic *C. sativa* expressing the crTCA cycle displayed significant alterations to their morphological phenotype and physiology when grown in a greenhouse. Earlier in their development, the crTCA plants were shorter, on average, than WT or EV counterparts but this difference decreased over time (**Fig 2, Supplementary Fig S6**). The transgenic plants ended their life cycle at the same height and time as the controls and did not show any significant changes in total yield (**Table 2**). Despite an observed reduced height phenotype, the crTCA lines had higher rates of net photoassimilation (A_net_) and a higher measure of stomatal conductance (g_s_) (**Table 2**). Though these increases in A_net_ at ambient CO_2_ concentrations were observed, no change in the A/Ci response was identified in any transgenic line expressing the crTCA cycle genes (**Supplementary Fig S7**). Based on chlorophyll fluorescence analysis, a slight reduction in the estimation of non-photochemical quenching (NPQ) was observed, but no other changes were found in terms of biochemical parameters, content or makeup (**Table 2**).

**Fig 2:**
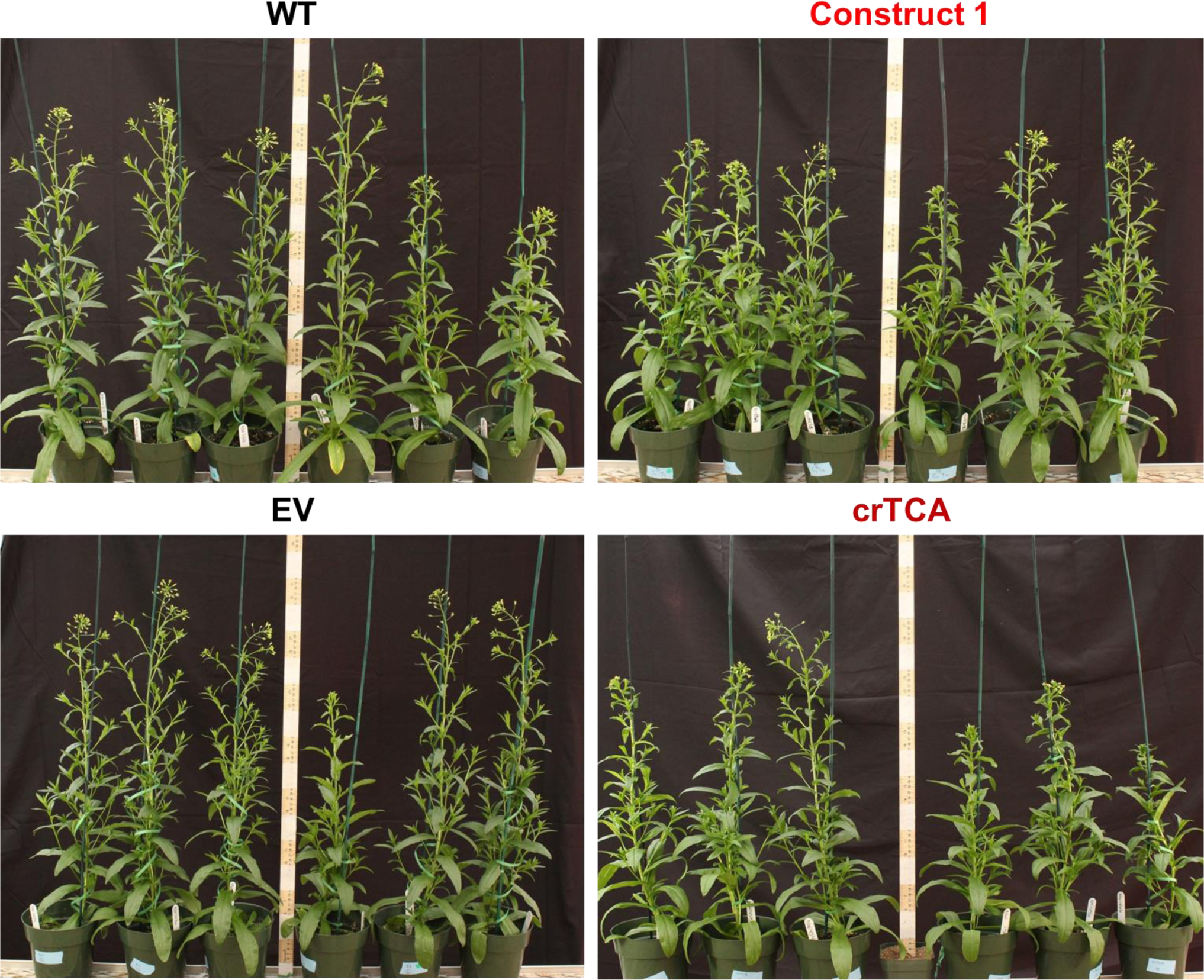
Phenotype of transgenic *C. sativa* grown in the greenhouse. Seven-week old plants expressing the crTCA cycle genes are noticeably shorter than non-segregant WT or Empty Vector controls. Construct 1 expression led to a slightly shorter plant height but also a noticeable increase in axillary branches.

**Table 2:** Gas exchange, chlorophyll fluorescence and seed yield of transgenic *C. sativa* grown in the greenhouse. Maximum rates of photoassimilation (A_net_) and stomatal conductance (g_s_) were analyzed on plants acclimated to a light intensity of 1200 µmol m^-2^ s^-1^ PPFD. Dark respiration (R_d_) and dark-adapted measurements of chlorophyll fluorescence was performed 2 hours before dawn. ΦPSII: quantum yield of PSII; F_v_/F_m_: maximum quantum yield of PSII; NPQ: non-photochemical quenching; Fv’/Fm’: light-adapted quantum yield of PSII. Means are ± the standard error of the mean (SEM), n ≥ 3 plants / line for each measurement.

|  | WT | EV | crTCA | Construct 1 |
| --- | --- | --- | --- | --- |
| $A_{\text{net}}$<br>($\mu\text{mol CO}_2 \text{ m}^{-2} \text{ s}^{-1}$) | $24.58 \pm 0.2^b$ | $24.12 \pm 0.3^b$ | **$27.58 \pm 0.2^a$** | **$18.53 \pm 0.2^c$** |
| $g_s$<br>($\mu\text{mol CO}_2 \text{ m}^{-2} \text{ s}^{-1}$) | $0.27 \pm 0.01^c$ | $0.30 \pm 0.01^b$ | **$0.39 \pm 0.01^a$** | $0.24 \pm 0.02^c$ |
| $R_d$<br>($\mu\text{mol CO}_2 \text{ m}^{-2} \text{ s}^{-1}$) | $-1.96 \pm 0.6^a$ | $-1.54 \pm 0.04^a$ | $-1.66 \pm 0.06^a$ | $-1.47 \pm 0.03^a$ |
| $\Phi\text{PSII}$ | $0.419 \pm 0.02^a$ | $0.424 \pm 0.02^a$ | $0.428 \pm 0.04^a$ | $0.410 \pm 0.08^a$ |
| $F_v/F_m$ | $0.813 \pm 0.005^a$ | $0.808 \pm 0.007^a$ | $0.799 \pm 0.016^a$ | $0.818 \pm 0.004^a$ |
| NPQ | $0.954 \pm 0.13^a$ | $1.03 \pm 0.05^a$ | $0.744 \pm 0.08^a$ | $1.15 \pm 0.0^a$ |
| $F_v'/F_m'$ | $0.587 \pm 0.02^a$ | $0.583 \pm 0.01^a$ | $0.618 \pm 0.01^a$ | $0.578 \pm 0.03^a$ |
| Chlorophyll a | $2.29 \pm 0.17^a$ | $2.02 \pm 0.07^a$ | $2.09 \pm 0.11^a$ | $2.24 \pm 0.05^a$ |
| Chlorophyll b | $0.601 \pm 0.06^a$ | $0.552 \pm 0.05^a$ | $0.548 \pm 0.04^a$ | $0.565 \pm 0.02^a$ |
| Total Chlorophyll | $2.90 \pm 0.23^a$ | $2.57 \pm 0.11^a$ | $2.26 \pm 0.15^a$ | $2.28 \pm 0.07^a$ |
| Total Yield<br>(g / plant) | $7.09 \pm 0.3^a$ | $6.69 \pm 0.3^a$ | $7.03 \pm 0.2^a$ | $7.37 \pm 0.3^a$ |
| Seed Weight<br>(mg / 50 seed) | $74.5 \pm 1.7^a$ | $73.9 \pm 1.2^a$ | $74.0 \pm 1.0^a$ | $78.8 \pm 0.8^a$ |

Because the crTCA genes were spread across multiple vectors, we also assessed these individual vectors or partial crTCA cycle lines. While these lines did not manifest as large of physiologic changes as the full crTCA cycle, we found that these vectors individually had unique outcomes on the phenotype and physiology of *C. sativa*. In the greenhouse, the expression of Construct 1 (OSR/ICDH, OGC and ICL) led to a decrease in net photosynthetic rate (A_net_) and in stomatal conductance but also resulted in a moderate increase in total seed yield, presumably due to the increased axillary branches in this line (**Fig 2, Table 2**).

RNA-Seq analysis found that the crTCA lines had little changes in their transcriptome with only 92 differentially expressed genes (DEGs, **Supplementary File S1**). Interestingly, partial cycle expression (e.g. Construct 1) led to greater changes in differential gene expression (n = 239) that reflected a stronger phenotype. To understand if certain transgenes of the crTCA cycle were correlated with specific transcripts we used weighted gene co-expression network analysis (WGCNA) [28]. From the lists of highly correlated genes, we used gene ontology (GO) to inform us about large-scale metabolic processes that are altered in the transgenic transcriptomes. The most significant GO terms that were found to be negatively correlated with crTCA expression were related to carboxylic- and oxo-acid metabolism. There was only one positive correlation found with crTCA expression with the GO term for amino acid biosynthesis (**Supplementary Table S6**)

### Elevated CO_2_ effects on transgenic *C. sativa* expressing the crTCA cycle

In the greenhouse, CO_2_ concentrations are approximately at atmospheric concentrations (∼415 ppm). We decided to grow the lines at elevated CO_2_ concentrations (1200 ppm) to test the effects of increased availability of CO_2_. We also tested two light intensities: 200 μmol m^−2^ s^−1^ and 1200 μmol m^−2^ s^−1^. Under the lower light intensity, the crTCA plants exhibited a similar phenotype as to that in the greenhouse. crTCA plants were significantly shorter than the EV control plants while Construct 1 plants had no discernible phenotype (**Fig 3A**). At the lower light intensity, crTCA plants had significantly increased rates of A_net_ and Construct 1 plants had significantly lower estimates of gs (**Fig 3B**). When these plants were grown at the same CO_2_ concentration but at a higher light intensity, the phenotype associated with the crTCA cycle disappeared but manifested in the Construct 1 plants (**Fig 3A**). In this environment, both transgenic lines had increased rates of A_net_ and larger estimates of gs and transpiration (**Fig 3B**). Construct 1 had higher overall rates of A_net_ while crTCA lines had higher estimates of stomatal conductance. Both transgenic lines had higher rates of dark respiration (R_d_) but no significant changes in yield or chlorophyll fluorescence parameters (**Supplementary Table S7**).

**Fig 3:**
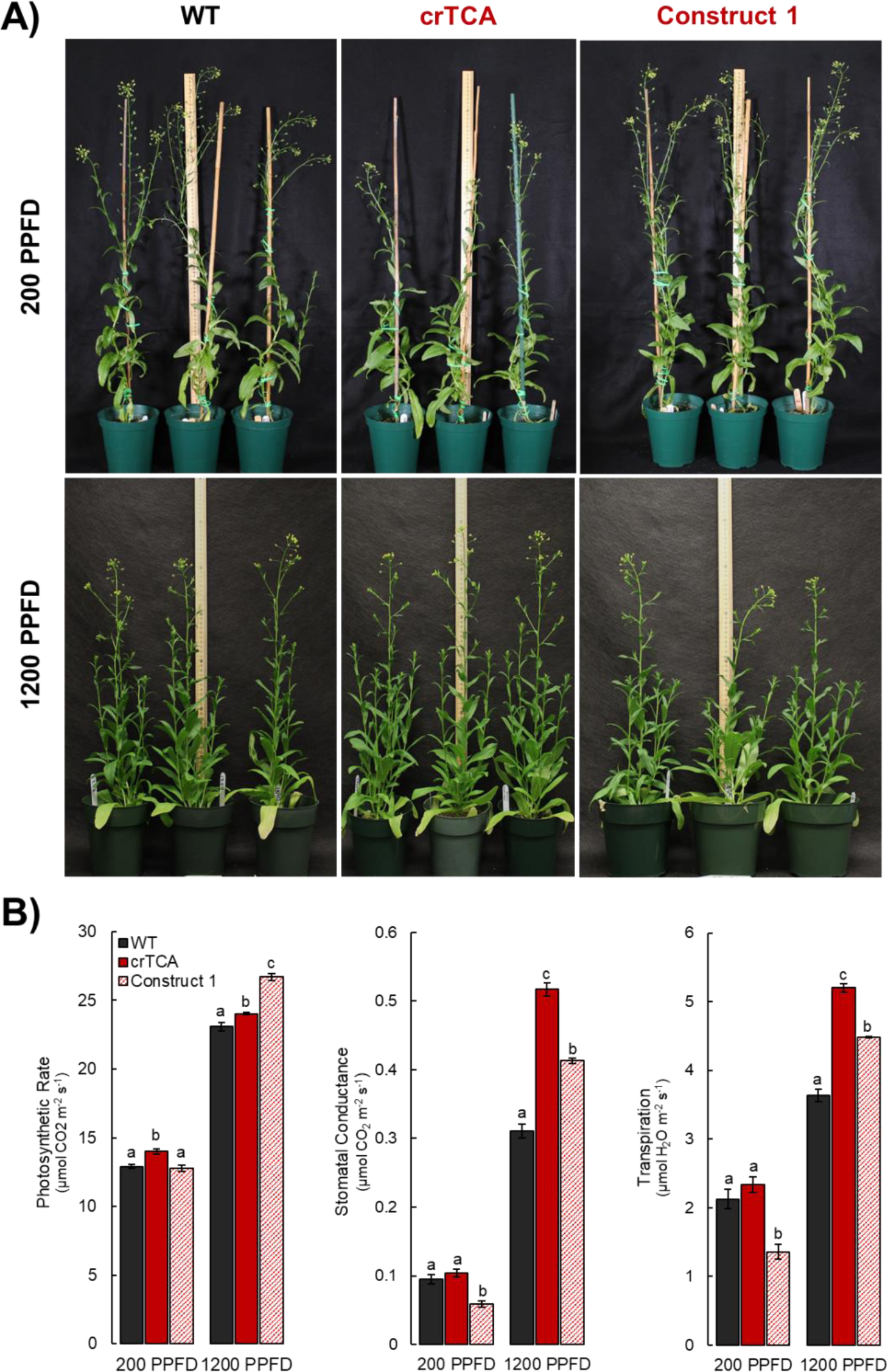
Light-dependent phenotype and physiology of transgenic *C. sativa* grown at elevated CO_2_ concentrations. A) At 1200 ppm CO_2_ and 200 PPFD (top row), plants expressing the crTCA cycle (middle column) appear shorter than both WT (left column) and the partial crTCA line, Construct 1 (right column). At the same CO_2_ concentration and 1200 PPFD (bottom row), plants expressing the crTCA cycle display no phenotype whereas Construct 1 plants are slightly smaller. B) Physiologic measurements of photosynthetic gas exchange parameters. n = 4 plants / line; letters indicate statistically significant groups determined via Tukey HSD, p < 0.05

### Metabolic changes of transgenic *C. sativa* grown at elevated CO_2_

Metabolic flux analysis was used to assess changes in central metabolism at elevated CO_2_ concentrations (600 ppm) and a saturating light intensity (1500 μmol m^−2^ s^−1^). The rate of both glycine and serine ^13^CO_2_ labelling is significantly reduced in transgenic crTCA plants (**Fig 4**).

**Fig 4:**
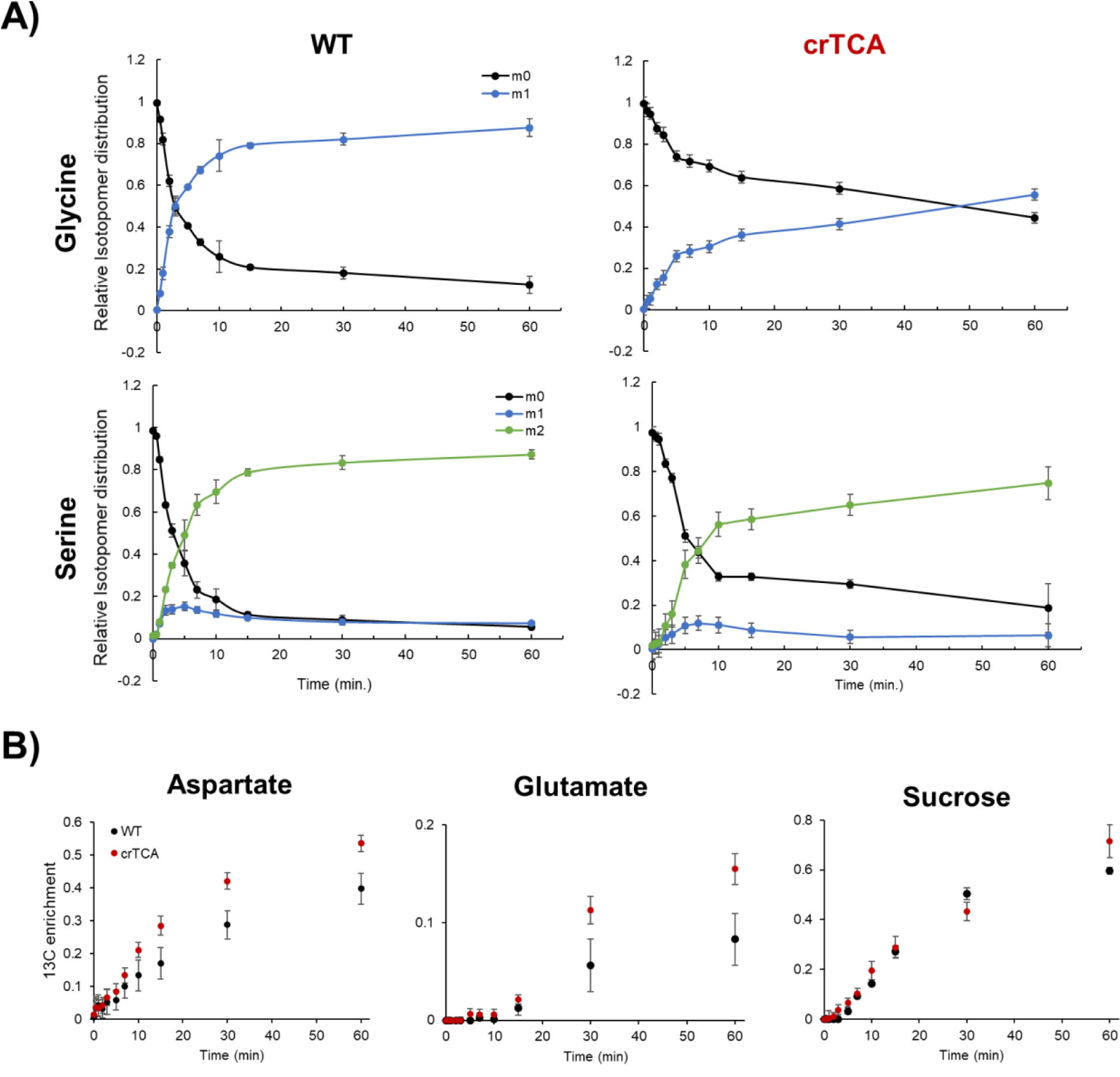
Transient ^13^CO_2_ labeling in glycine and serine in WT and crTCA lines. A) The mass isotopomer distribution (MID) of glycine and serine is shown where the points represent averages and error bars represent ± one standard deviation. Glycine is the top row, serine is the bottom. WT is the left column and crTCA is the right column. For each time point, n = 3 biological replicates. Each color represents a distinct isotopomer with various degrees of ^13^C incorporation: m0 is unlabeled, m1 is one ^13^C label; m2 is two ^13^C labels. B) ^13^C enrichment of aspartate, glutamate and sucrose. WT samples are the black circles while crTCA samples are the red circles. The mean is presented with error bars representing one standard deviation (n = 3).

For both glycine and serine, the fraction of molecules that are unlabeled (giving the isotopomer m0) quickly decreases while the fraction of labeled isotopomers (m1 or m2) rises rapidly in the WT samples. In the crTCA lines, the labeling occurs at much slower rates. In the crTCA line, glycine labelling reaches a significantly slower level, with only about 50% of the total glycine pool being labelled after ∼60 minutes whereas in WT 50% of the total glycine pool is labelled within ∼5 minutes. Serine was fractionally labelled slightly faster than glycine, however, the rate of the crTCA plants is, again, significantly slower than WT. Aspartate and glutamate, however, showed the opposite trend – with greater ^13^C enrichment in the crTCA lines than in the WT (**Fig4B**). Notably, many of the central metabolism intermediates did not show any differences between the WT and crTCA lines in the time period assayed. Several metabolites related to the TCA cycle (e.g. malate, succinate, fumarate and citrate) show very limited labeling within the 60 minutes given for this experiment and no changes were detected between WT and transgenic lines (**Supplementary Fig S9**). A preliminary analysis of steady-state levels only revealed significant differences in pyruvate and fructose in both crTCA and Construct 1 lines (**Supplementary Table S9**).

## Discussion

We present the first iteration of a build, test, and learn cycle [29] of a short synthetic carbon fixation cycle that was inspired by theoretical consideration of such a synthetic metabolism [18]. This is the first demonstration of a reverse TCA cycle-based synthetic carbon fixation cycle *in vitro*, and the first demonstration of a functional four-step carbon fixation cycle. The condensed, reverse TCA (crTCA) cycle described in this study has been shown to fix carbon *in vitro* and the presented enzymes retain activity when expressed *in planta*. When stably transformed into *C. sativa*, we found that a core enzyme of the cycle, KOR, had undetectable protein levels despite a high level of expression (**Supplementary Fig S5, Supplementary Table S5**). It is therefore unlikely that the crTCA “cycle” is operational in *Camelina*. However, we found that the presence of a partial crTCA cycle or partial crTCA pathway had a substantial impact on the morphological phenotype, physiology and metabolism of *Camelina*.

The enzymes of the crTCA cycle are not found in any one organism and therefore each enzyme hails from different species of bacteria (**Table 1**). Several candidate enzymes were screened for each crTCA cycle step and the best enzyme from these candidates was chosen. The five enzymes of the crTCA cycle were evaluated *in vitro* for their activity and we moved forward with *in planta* expression. Unfortunately, we found that our chosen OGC enzyme had no or little activity *in planta* as an oxoglutarate carboxylase. However, we also found that our chosen OSR enzyme functions as an isocitrate dehydrogenase (ICDH), capable of performing both the carboxylation of 2-oxoglutarate and the subsequent reduction of oxalosuccinate to isocitrate (**Fig 1, Table 1**). While initially designed to function with five enzymatic steps, it was demonstrated that this version of the cycle could function with only four steps. This does not mean that a five-step crTCA cycle is not needed, but suggests that careful enzyme assessment and selection is required, particularly in identifying an OGC enzyme. In the initial description of a shortened, reverse TCA cycle, the authors raise concerns about the function of this cycle under physiologic conditions due to thermodynamic bottlenecks [18]. These concerns are somewhat assuaged by the inclusion of a fifth enzyme (2-oxoglutarate carboxylase, OGC), but our choice for this enzyme had low activity when expressed *in planta* and may have limited the effects of crTCA expression (**Table 1**).

Another concern on the feasibility of the crTCA cycle is the ability of the cycle to operate in the reverse direction. This reverse reaction was demonstrated *in vitro* in this study. When the crTCA cycle is provided with glyoxylate and succinate as starting material, the amount of glyoxylate decreases rapidly (**Supplementary Fig S1**). This creates a “see-saw” effect that limits the accumulation of multiple ^13^C-labeled metabolites, especially *in vitro* and may be an artifact of our assay. This potential for reversibility is less understood *in planta*. When expressed *in planta*, we hypothesize that reversibility will be less of an issue as glyoxylate metabolism is a robust system that is operational in the plastid and peroxisome [30, 31]. Likely substrates for the crTCA that are present in the plastid are succinate, 2-oxoglutarate and isocitrate. Such dicarboxylates have been shown to cross freely past the outer membrane of the chloroplast and have bidirectional transport through the inner membrane to the stroma [32]. All three can be present in the cytosol as well. Succinate is primarily found in the mitochondria, where it is involved in the TCA cycle, but can also be found in the peroxisome and cytosol where it participates in both the glyoxylate cycle and GABA shunt respectively [33]. Succinate has also been identified in the chloroplast, but at a lower concentration: approximately 3.36 nmol g FW^-1^ in the plastid and 14.3 nmol g FW^-1^ in the cytosol [32, 34]. 2-oxoglutarate is present in the plastid where it plays a central role in amino acid biosynthesis and ammonia assimilation, being the substrate for glutamate via GOGAT. In the leaves of *A. thaliana,* the concentration of 2-oxoglutarate is approximately 14.3 nmol per gram fresh weight (FW) in the plastid and 22.1 nmol/g FW^30^. Isocitrate is intricately linked to 2-OG, being its precursor in both the mitochondria and the cytosol [33].

The crTCA cycle uses enzymes from the host organism without modification. The enzymes of the crTCA cycle were sourced from a suite of microbial organisms from diverse environments. This study expressed these enzymes in the context of the chloroplast - which is a dynamic environment unlike their host microbes. The activity and directionality of the crTCA cycle may change depending on environmental parameters. For instance, the availability of light changes the stromal pH (pH 8 in the light; and pH 7 in the dark) and the plastidial concentration of ATP and NADH are reduced by a factor of ∼2.4 at night [36]. The crTCA cycle’s activity was measured *in vitro* at pH 8.0 and was provided with excess ATP, NAD(P)H, and substrate (see Methods). Rational enzyme engineering approaches are capable of using site-directed mutagenesis to enhance enzymatic activity for their desired purposes [37–39]. Possibilities for increasing crTCA enzyme efficiency would be to increase abundance and activity (OGC and KOR are prime targets), reduce reversibility (ICL) or to design the enzymes to better suit the environment of the plant chloroplast (i.e. pH changes, temperature). This has been done in other *in vitro* carbon fixation cycles [20], but requires an abundance of information on the enzymes in question (e.g. structural information) that is not readily available for any crTCA enzyme. Enzyme engineering is crucial for the development of subsequent iterations of this synthetic cycle. These approaches would provide additional *in vitro* support for the function of the crTCA cycle when expressed *in planta*.

Despite the room for improvement, the first iteration of the crTCA cycle did allow us to prove that this metabolism can increase net CO_2_ assimilation in *C. sativa*. In both a greenhouse environment and when grown under elevated CO_2_, the transgenic plants had increased rates of photosynthetic gas exchange (**Tables 2, 3**). The correlation of A_net_ with changes in gs and transpiration are indicative of a stomatal mechanism rather than a carboxylation mechanism. Initially, we hypothesized that malate/fumarate levels [33,40,41] could be altered in crTCA lines due to potential metabolic interaction with the mitochondrial TCA cycle, however in our preliminary metabolic analysis, no significant changes were detected in sampled mesophyll tissues (**Supplementary Table S8**). Alternative mediators of stomatal opening and closure, such as ABA or H2O2 may be involved but were not investigated in this study. Furthermore, in no environment was a positive A/Ci response detected (**Supplementary Fig S6**). Interestingly, several lines of evidence suggest the photosynthetic changes of the transgenic *C. sativa* was found to be more closely associated with light intensity than with CO_2_ concentration. This is interesting to consider with the morphological changes seen in the crTCA lines, with the predominant phenotype of the crTCA lines being shorter overall than WT or EV controls. Only when grown in high CO_2_ and high light intensities were crTCA plants the same height as controls (**Figs 2, 3**).

Glyoxylate is known to be present in photorespiration and is the downstream product of glycolate oxidation in the peroxisome [31]. Glyoxylate can then transaminated by the enzyme GGAT (glutamate:glyoxylate aminotransferase) to yield glycine. GGATs can use either glutamate or alanine as the amino donor to yield 2-oxoglutarate or pyruvate, respectively.

Glutamate showed a higher rate of ^13^C enrichment without changing its relative pool size (**Fig 4B**). Steady-state levels of pyruvate were found to be significantly higher and alanine may also be increased but additional replicates are required (**Supplemental Table S9**). Overexpression of the GGAT enzymes in *A. thaliana* has led to increases in both steady-state levels of serine and glycine content in leaf tissue [45]. Whereas knockout studies in GGATs have shown to have a reduced phenotype and also are altered ABA and H2O2 metabolism [45–47]. Glyoxylate can additionally be transaminated by SGAT (serine:glyoxylate aminotransferase) to yield glycine. The SGAT reaction utilizes serine as amino donor to yield hydroxypyruvate which can be reduced to glycerate for re-introduction into the CBC. Studies with increased SGAT activity have shown reduction in plant biomass associated with increased serine content at the expense of carbohydrate and sugar contents [48, 49]. Additional metabolic investigation into serine and alanine contents will greatly advance our understanding of the crTCA cycle and its contribution(s) to central metabolism and photorespiration.

Photorespiration is seen as an energetically costly process, with estimates of energy loss ranging from 30-50%. Despite these costs, recent reexaminations of photorespiration suggest that the amino acids glycine and serine may serve as an additional carbon sink of plant metabolism [50]. While more work is required, amino acid pools may represent the carbon sink for additionally fixed carbon via the crTCA cycle. This may be an advantageous trait in the context of C/N balancing, especially in environments of elevated CO_2_. It is well documented that several crop species have reduced N assimilation when grown at elevated CO_2_ and this directly affects yield potential. Additionally, lower N assimilation at elevated CO_2_ results in reduced photosynthetic investment with both RuBisCO and/or total protein content and chlorophyll content decreasing [51, 52]. This phenomenon therefore may be a substantial risk to food security in the next century as both CO_2_ concentrations and population growth continue to rise [51,53– 55]. While photorespiration is often thought to be a wasteful, energy-consuming process, there is growing support that the reactions also work to balance and re-distribute energy necessary for optimal photosynthesis, especially in the context of elevated CO_2_ [50,54,56,57].

In summary, we present the first iteration of a build-test-learn cycle of a condensed, reverse TCA (crTCA) cycle both *in vitro* and *in planta*. This study realizes the design of an *in vitro* carbon fixation cycle initially posited by Bar-Even et al. [18]. We demonstrate that this simple metabolism can fix carbon *in vitro* and change carbon metabolism when expressed *in planta*. The mechanism behind this increase is not fully known but appears to be related to changed stomatal behavior and/or changed photorespiratory metabolism. Future metabolite profiling and flux measurements are crucial to understanding and defining the effects of the crTCA cycle. Furthermore, engineering the crTCA enzymes for better performance *in planta* is also imperative for future version of this metabolism.

## Materials & Methods

### Selection of crTCA Enzyme Candidates

To generate a robust gene candidate list (3-5 target genes) for each of the five enzyme steps of the proposed synthetic crTCA cycle, BLAST-p (NCBI) alignments and Conserved Domain (NCBI) analysis were used to identify target crTCA cycle enzymes and appropriate catalytic/substrate binding domains. The query sequences for the BLAST-p alignments were from enzymes known to have activity for crTCA cycle functions [21, 58]. To identify suitable enzymes for each catalytic step, we used blastp with the following query sequences: Succinyl-CoA synthetase (*E. coli* [59]; NP_415257.1); 2-oxoglutarate : ferredoxin oxidoreductases (*Hydrogenobacter thermophilus*; alpha-SU YP_003433044; beta SU WP_012963730.1) 2-Oxoglutarate Carboxylase (*Hydrogenobacter thermophilus*; SSU YP_003433054.1; LSU, YP_003433044); Oxalosuccinate Reductase (*Chlorobium limicola*; BAC_00856.1); Isocitrate Lyase (*Corynebacterium glutamicum;* NP_601530.1) [21,25,26,60–62]. MODELLER [63] was used to predict the binding affinity of the candidates for the cycle substrates.

The selected genes were synthesized by GenScript with codon optimization for *E. coli* and ligated into pUC57. Enzymes 1-3 are multi-subunit, and the sequences for these enzymes were synthesized consistent with the NCBI genome sequence for each organism including intergenic spacer regions, which were not codon optimized. Ferredoxin from *Hydrogenobacter thermophilus* TK-6 was also synthesized to supplement the KOR enzyme in the proposed crTCA cycle. The selected candidates are listed (**Table 1**).

### In vitro expression and purification

Qiagen pQE vector system was utilized for the overexpression of the crTCA cycle enzymes with of His-tags for downstream purification. Some of the crTCA enzymes did not express well with the pQE vector and were instead expressed using pET21b or pET28a (MilliporeSigma) as indicated in **Supplementary Table S8**.

The primers and annealing temperatures used to amplify the candidate crTCA cycle genes are also listed in **Supplementary Table S8**. For cloning the genes into pQE-1, forward primers were designed to amplify the sequence beginning with the 5’ ATG to limit additional amino acids on the N-terminus. Reverse primers included the *Hind*III or *Sac*I sites of pUC57. The synthesized crTCA cycle genes in pUC57 were used as template DNA with iProof High-Fidelity DNA polymerase (BioRad). The PCR products were gel purified, digested with *Hind*III or *Sac*I, and precipitated with ethanol. Following phosphorylation with T4 polynucleotide kinase (New England BioLabs), the PCR products were ligated into expression vector pQE-1 (Qiagen) with T4 ligase (New England BioLabs), and the ligation mix was transformed into *E. coli* strain XL-1 Blue. Plasmid DNA was confirmed by sequence and then transformed into expression strain *E. coli* M15. Genes for BrBT-SCS and HyTh-KOR were also cloned into pET21b and pET28a, respectively. PCR products were ligated into pPCRscript and transformed into *E. coli* XL-1 Blue. Digestion with *Nde*I and *Xho*I was conducted to extract the PCR product from pPCRscript using sites in the PCR primers. The digested PCR product was gel extracted, ligated into pET21b or pET28a and transformed into *E.coli* XL-1 Blue. Plasmid DNA was confirmed by sequencing before transformation into expression strain *E.coli* BL21 (DE3).

The crTCA cycle candidate proteins were expressed as N-terminal his-tag fusions, purified and evaluated for activity. Enzymes with the highest activity were selected for LC-MS assays and transient expression in tobacco. BrBT-SCS was expressed in *E. coli* BL21 (DE3) transformed with BrBT-SCS in pET21b. The cells, grown in LB at 37°C and 200 rpm to mid-log phase (OD_600_ 0.6-0.8), were induced with 0.1 mM IPTG and the temperature was reduced to 18°C for 18-20 h. HyTh-KOR was expressed in *E. coli* BL21 (DE3) transformed with HyTh-KOR in pET28a. The cells, grown in LB, supplemented with 1 mM thiamine, at 37°C and 200 rpm to mid-log phase (OD_600_ 0.6-0.8), were then induced with 0.1 mM IPTG. FeSO4 (0.5 mM) was added to increase iron availability for Fe-S cluster biosynthesis. The temperature was then reduced to 18°C for 18-20 h. MaFe-OGC was expressed in *E. coli* M15 transformed with MaFe-OGC in pQE1. Cells were grown in LB, with 1 mg/L biotin, at 25°C, 200 rpm to OD_600_ 0.6-0.8, and expression was induced with 0.05 mM IPTG. The temperature was reduced to 18°C and incubated for an additional 18-20 h at 200 rpm. For NiHa-OSR and NoFa-ICL enzymes, freshly transformed *E. coli* M15 cultures were grown at 25°C, 200 rpm, to mid log phase (OD_600_ 0.6 to 0.8). IPTG was added (to 0.05 mM) and cultures were shaken at 200 rpm and 15°C, for 16 to 18 h. For all enzymes, prior to increasing the cultures to 1 L, soluble protein expression was confirmed from a 30 ml culture using SDS-PAGE and/or Western blot analysis.

### Purification of Recombinant crTCA Cycle Enzymes

All purifications used a BioRad DuoFlow system. Cell pellets containing the recombinant crTCA cycle proteins were resuspended in 50 mM sodium phosphate, pH 7.5, containing 1 mM benzamidine–HCl. All buffers for the KOR enzymes contained 0.01% Triton X100 during the purification [60]. The bacteria were passed through a French pressure cell (1,100 lb per in^2^) two to three times. The lysate was centrifuged at 15,000 x g for 40 min at 4°C to remove cell debris, then passed through a 0.45 µm syringe filter. The filtrate was applied to a 5 mL HisTrap HP Nickel Sepharose™ affinity column (GE Healthcare Life Sciences) and washed with five column volumes of binding buffer (50 mM sodium phosphate buffer, 500 mM NaCl, 30 mM imidazole, pH 7.5). The elution buffer was 50 mM sodium phosphate, 300 mM NaCl, 250 mM imidazole, pH 7.5. Elution was done via a linear gradient from 0% to 100% elution buffer. Fractions were analyzed by SDS-PAGE (12.5%) and protein concentration was estimated using the BioRad Bradford Assay (with a BSA standard). Fractions containing recombinant protein were pooled prior to additional purification. *BrBT-SCS:* Pooled fractions were dialyzed using (20 kDa MWCO) Slide-A-Lyzer (Thermo Fisher Scientific), in 50mM Tris-HCl (pH 8) overnight at 4°C. Dialyzed protein was then loaded onto a 5 ml HiTrap Q HP anion exchange column (GE Healthcare Life Sciences) and eluted with a gradient of 0-100% elution buffer (50 mM Tris-HCl (pH 8), 1 M NaCl). Fractions containing BrBT-SCS were dialyzed again as described, quantified, and stored at -80°C in 15% glycerol. *HyTh-KOR:* Microaerobic handling was required for the KOR enzymes, which involved de-gassing and sparging of all purification buffers with argon and anaerobic collection of elution fractions. After affinity chromatography, HyTh-KOR was desalted into 20 mM Tris-HCl (pH 8), 0.01% Triton X100 using a 30 kDa MWCO centrifugation filter (MilliporeSigma) in an anaerobic glovebox. Purified enzymes were quantified, and stored anaerobically at -80°C in stoppered vials in 15% glycerol. *MaFe-OGC:* Pooled fractions were dialyzed using Slide-A-Lyzer (ThermoFisher Scientific) cassettes (30 kDa MWCO) against 50 mM Tris-HCl (pH 8) then stored at -80°C in 15% glycerol. *NiHa-OSR and NoFa-ICL:* Fractions containing recombinant protein were pooled and dialyzed using a Slide-A-Lyzer (ThermoFisher Scientific) cassette (20 kDa MWCO) against 50 mM Tris-HCl, 1 mM benzamidine, 0.25 mM EDTA, pH 8.0. The dialyzed samples were applied to a 5 ml HiTrap Q HP anion exchange column (GE Healthcare Life Sciences). The Q anion exchange column was eluted via linear gradients from 0% to 100% elution buffer (50 mM Tris, 1 M NaCl, 1 mM benzamidine, 0.5 mM EDTA, pH 8). Appropriate fractions were pooled and dialyzed using a Slide-A-Lyzer (Thermos Fisher Scientific) cassette (20 kDa MWCO) against 50 mM Tris-HCl, 1mM benzamidine, 0.25 mM EDTA, pH 8.0. The purified enzymes were stored at -80°C. in 15% glycerol. *HyTh-FDX:* Fractions were pooled and loaded onto a HiPrep 26/10 desalting column (GE Life Sciences) into 50 mM Tris-HCl, pH 8. Protein was stored at -80°C in 15% glycerol.

### Spectrophotometric and LC-MS Assays for crTCA Cycle Enzymes

Spectrophotometric assays were carried out to screen the candidate enzymes for each crTCA cycle step *in vitro* as well as to detect *in planta* activity in tobacco. A Biomate 3 spectrophotometer from ThermoFisher Scientific and a Shimadzu UV-2401PC UV-visible spectrophotometer with a temperature controlled cuvette holder were used for the assays. In all cases, individual enzyme reactions were conducted in triplicate using purified enzymes from the same batch of purification that had been stored in aliquots at -80°C. *SCS:* The standard reaction consisted of 10 mM sodium succinate, 10 mM MgCl_2_, 0.1 mM CoA, 0.1 mM DTT, 0.4 mM nucleotide ATP and 0.1 M KCl in 50 mM Tris-HCl (pH 7.4). Reactions were started with purified enzyme or extracts of cells transformed with BrBT-SCS. The reaction was monitored by absorbance at 230 nm in response to thioester formation at RT [64]. *KOR:* The decarboxylase activity of KOR was detected in a continuous spectrophotometric assay following the enzyme-, substrate-, and time-dependent reduction of oxidized benzyl viologen (at 600 nm). Reaction mixtures were prepared aerobically, then sparged with argon. The assay was performed in anaerobic gas-tight glass cuvettes. The KOR enzyme was treated with 5 mM DTT (15 min) prior to adding the enzyme to the reaction mix. The reaction mix contained 100 mM sodium phosphate (pH 7.5), 1 mM benzyl viologen, 2.5 mM 2-OG, 0.5 mM coenzyme A, 4 mM MgCl_2_ and 0.025 mM sodium dithionite. The reactions were started by adding 2-OG using gas-tight glass syringes. The assay was conducted at RT or 30°C. *OGC:* A two-step, coupled spectrophotometric assay was developed for the ATPase activity of OGC using phosphoenolpyruvate kinase (PK) and lactate dehydrogenase (LDH). PK utilizes ADP hydrolyzed by OGC to produce pyruvate from phosphoenolpyruvate, which is converted to lactate by LDH by the oxidation of NADH. The oxidation of NADH is observed by absorbance at 340 nm. The first step reaction mixture is composed of 100 mM PIPES (pH 6.5), 5 mM MgCl_2_, 20 mM 2-OG, 50 mM NaHCO_3_, and 5 mM ATP. The reaction was initiated by OGC. The reaction mixture with OGC was held for 30 min at RT (65°C for the thermophilic enzyme). For the second step, 0.1 mM β-NADH, 2 mM phosphoenolpyruvate and PK/LDH were added to the first step reaction, and NADH oxidation was monitored by absorbance at 340 nm. The amount of ADP produced was estimated using a standard curve. *OSR:* The assay evaluates the dehydrogenase activity of OSR, monitored at 340 nm, measuring the reduction of NADP+. The reaction mixture is composed of 50 mM Tris (pH 7.4), 10 mM MgCl_2_, 100 mM KCl, 4 mM isocitrate, 4 mM β-NADP^+^. The reaction was initiated by addition of purified enzymes or tobacco cell extracts and monitored by NADP^+^ reduction (340 nm at RT). The OSR carboxylation assay was adapted from a published method [26]. The reaction mixture contained 50 mM PIPES (pH 6.5), 10 mM 2-OG, 1 mM NADH, 10 mM MgCl_2_, 50mM NaHCO_3_. The reaction was started with the addition of the NiHa OSR enzyme. The activity is measured following the oxidation of NADH (340 nm at RT). *NoFa ICL:* Reaction mixtures contained 30 mM imidazole (pH 6.8), 5 mM MgCl_2_, 1 mM EDTA, 4 mM phenylhydrazine and 10 mM isocitrate. The reaction was performed at RT by adding purified protein or cell extracts and monitoring the absorbance of glyoxylate phenylhydrazone (324 nm) in the presence of phenylhydrazine [62]. *Pyruvate Carboxylase:* This assay evaluates pyruvate carboxylase activity of MaFe OGC. It is a coupled assay, where oxaloacetate is first produced by pyruvate carboxylase. The oxaloacetate is then used by malate dehydrogenase to produce malate by the oxidation of NADH, measured by absorbance at 340 nm [65]. Reaction mixtures contained 50 mM Tris-HCl (pH 8.0), 8 mM MgCl_2_, 20 mM sodium pyruvate, 20 mM NaHCO_3_, 8 mM ATP, 0.2 mM NADH, and 50 μM acetyl-CoA. The reaction was started by addition of 1 U of porcine heart malate dehydrogenase (MilliporeSigma) and MaFe OGC. The production of NAD^+^ was measured by absorbance at 340 nm at RT.

High performance liquid chromatography-mass spectrometry (HPLC-MS) analysis assays were developed for those enzymes that could not be assayed spectrophotometrically for the direction of the crTCA cycle. Electrospray ionization-mass spectrometry (ESI-MS) analyses were performed on an Agilent LC-MS system comprised of an Agilent 1200 series HPLC with an Agilent 1260 Infinity micro degasser, binary pump, and standard auto-sampler, an Agilent 1290 Infinity diode-array detector and an Agilent 6520 Accurate-Mass Q-TOF spectrometer, equipped with an electrospray ionization interface. The enzyme reaction samples were separated using a modified version of a published method [66]. A Synergy Hydro-RP column (100 mm × 2 mm, 2.5 μm particle size, Phenomenex), was used for reversed phase chromatography. The total run time is 18 min with a flow rate of 0.200 ml/min. All solvents used for LC-MS were LC-MS grade. Solvent A was 0.1% formic acid in water; solvent B 0.1% formic acid in methanol. The gradient used was 0 min, 0% B; 10 min, 60% B; 11 min, 0% B; 17 min, 0% B. The ESI-MS was set in the negative ion mode with spectra acquired over a mass range from m/z 50 to 1000. The optimum values of the ESI-MS parameters were: capillary voltage, +3.5 kV; liquid nitrogen used as dry gas; drying gas temperature, 325 °C; drying gas flow rate, 10.0 L/min; nebulizing gas (pure nitrogen gas) pressure, 35 psi; fragmentor voltage, 115V. Reactions were stopped by addition of 50 µl methanol and processed by LC-MS as described above, unless otherwise stated.

*OGC/OSR (step 3-4):* The OGC product, oxalosuccinate is labile, so the production of isocitrate for the OGC/OSR coupled reaction was tested by LC-MS. The reaction contained 50 mM Tris (pH 6.5), 5 mM MgCl_2_, 20 mM 2-OG, 50 mM NH4HCO_3_, 5 mM ATP, 50 mM KCl, 2 mM β-NADPH and recombinant OGC and OSR enzymes. The reaction was initiated by adding the enzymes and held for 30 min at RT. The isocitrate product was analyzed by LC-MS.

#### Partial Function of crTCA Cycle (step 3-5)

The reaction contained 50 mM Tris-HCl (pH 6.5), 5 mM MgCl_2_, 20 mM 2-OG, 50 mM NH4H^13^CO_3_, 5 mM ATP, 50 mM KCl, 2 mM β-NADPH and the recombinant OGC, OSR and ICL enzymes. The reaction was initiated by adding enzyme and held for 30 min at RT. The final products, glyoxylate and succinate, and the intermediate product, isocitrate were identified by LC-MS.

#### Full Function of crTCA Cycle

To demonstrate *in vitro* function of the crTCA cycle, LC-MS samples were prepared (under anaerobic conditions) for all five crTCA cycle enzymes. In addition to the crTCA cycle enzymes, the reactions contained 50 mM Tris-HCl (pH 8.0), 4 mM NADH, 5 mM ATP, 5 mM MgCl_2_, 10 mM NaH^13^CO_3_, 0.5 mM CoA, 100 µM succinyl-CoA, 1 mM 2-OG, and 60 µg HyTh ferredoxin, in 600 µL. The reactions were initiated by addition of HyTh KOR and HyTh Fd, and incubated for 0, 15, 30, 60, 90 and 120 minutes at 30°C. The ^13^C-labeled crTCA cycle intermediates were analyzed by LC-MS, and quantification of the different labeled species of the crTCA cycle intermediates was performed using standards analyzed alongside the experimental samples.

#### crTCA Cycle Function Under Aerobic Conditions

The *in vitro* crTCA cycle reactions were prepared (under aerobic conditions) to test cycle function when the crTCA cycle enzymes were exposed to air. The aerobic reactions were prepared on the bench top instead of in an anaerobic glovebox.

#### The Reversibility of the crTCA Cycle

To test whether the crTCA cycle is reversible, the reactions contained 50 mM Tris-HCl (pH 8.0), 4 mM NAD^+^, 5 mM ADP, 5 mM MgCl_2_, 0.5 mM CoA, 500 µM succinate, 500 µM glyoxylate, and 60 µg HyTh ferredoxin, in 600 µl. The samples were prepared under anaerobic conditions. The reactions were initiated by addition of HyTh KOR and HyTh Fdx, and incubated for 60 minutes at 30°C. To further assess the reversibility of the ICL and OSR enzymes, a coupled assay was adapted from previous work [67]. The assay couples ICL with OSR catalyzing the conversion of glyoxylate and succinate to isocitrate by ICL, and finally to 2-OG through the activity of OSR with the oxidation of NAD^+^. The reactions contained 50 mM Tris-HCl (pH 8), 5 mM MgCl_2_, 1 mM NAD^+^, 0.5 mM glyoxylate, and 0.5 mM succinate. The reaction was started with the addition of ICL and OSR in a 1:1 ratio and was monitored for NADH using a Shimadzu UV-2401PC UV-Visible spectrophotometer (340 nm).

### Transient Expression in Tobacco

To validate the expression and activity of the crTCA cycle enzymes in a plant, each enzyme was transiently expressed individually in tobacco. The selected crTCA cycle gene sequences for each step were synthesized using GenScript with codon optimization for *Camelina sativa*. Each crTCA cycle gene was linked to a cauliflower mosaic virus (CaMV) 35S promoter, a chloroplast localization sequence (from tobacco), and the nopaline synthase (NOS) terminator. 6X-His tags were fused to the N-terminus for NiHa OSR and NoFa ICL and the C-terminus for the BrBT SCS and HyTh KOR coding sequences for the detection and purification of the proteins. The inserts for each crTCA cycle enzyme were ligated to the multiple cloning sites of pCAMBIA-EGFP [68] using *Hind*III and *Bam*HI sites, and transformed into *E. coli* XL1-Blue. The clones were screened using restriction analysis and verified by sequencing (Eurofins Genomics). The verified constructs were transformed into competent *Agrobacterium tumefaciens* GV3101 by electroporation and selected using kanamycin.

Each construct was transformed into the leaves of 5-6 week old tobacco plants using *Agrobacterium* infiltration [69]. BrBT-SCS, NiHa-OSR, and NoFa-ICL were all expressed in *Nicotiana tabacum*, while HyTh-KOR was expressed in *N. benthamiana* as expression with *N. tabacum* did not yield detectable protein. *A. tumefaciens* GV3101 harboring each crTCA cycle construct was grown in liquid yeast extract peptone (YEP) media with kanamycin, rifampicin, and gentamicin at 28°C until mid-log phase (OD_600_ 0.6-0.8). The cultures were centrifuged at 4,500 x g for 30 min. The pellets were resuspended in sterile infiltration media (10 mM MES, 10 mM MgCl_2_, 0.5% glucose, 200 µM acetosyringone, at pH 5.6) to an OD_600_ between 0.2-0.5 in 1 L of infiltration media. Containers of *Agrobacterium* suspension media were placed inside a desiccator. The leaves of each plant were submerged in the suspension by placing the plant upside-down. For the infiltrations with *N. benthamiana*, additional *Agrobacterium* GV3101 cultures were prepared containing the p19 viral suppression plasmid [70] and a 1:1 mixture of p19 culture to crTCA construct culture were added to the vacuum chamber. A vacuum of approximately 25 mm Hg was applied for 30 sec and released. The vacuum procedure was repeated for 3 cycles, after which the plants were allowed to dry and returned to the light shelf. The *N. tabacum* plants were grown with 12 h light and 12 h dark at 25°C, while *N. benthamiana* were grown with 16 h light (150 µmol m^-2^ s^-1^) and 8 h dark at 25°C.

Tobacco leaves were harvested 4 days post-transformation and the tissues were ground in liquid nitrogen by mortar and pestle. Proteins were extracted from ground tissue in a buffer containing 50 mM Tris (pH 8), 150 mM NaCl, 10% (v/v) glycerol, 1% (v/v) Triton x-100, and 1:100 protease inhibitor cocktail for plant cells (MilliporeSigma). Either crude tissue lysate or Ni-NTA Spin Kit (Qiagen) was used to prepare samples for Western blots. The extracts were mixed with Laemmli buffer containing 2-mercaptoethanol, boiled for 10 min, and fractionated by 12.5% SDS-PAGE. Proteins were transferred to a PVDF membrane (BioRad) using a Trans-blot Turbo System (BioRad). The membranes were blocked with 5% (w/v) non-fat dry milk in Tris-buffered saline and 0.1% (v/v) TWEEN 20 (TBST) overnight. The primary antibody used for BrBT-SCS, NiHa-OSR, and NoFa-ICL was the Penta-His antibody (Qiagen) at 1:4000 in TBST. Horseradish peroxidase (HRP)-conjugated goat anti-mouse (Seracare) was used as the secondary antibody at 1:8,500. For the HyTh-KOR, the primary antibody used was a polyclonal antibody raised against a HyTh-KOR specific peptide epitope. The primary antibody was diluted 1:5,000 in TBST with 1% (v/v) casein (MilliporeSigma). The secondary antibody was the HRP-conjugated goat anti-rabbit (Seracare) diluted at 1:20,000 in TBST with 2.5% (w/v) dry milk. Blot immunoreactivity was visualized by Clarity Western ECL Substrate (BioRad) and exposure to X-ray film.

To demonstrate the activities of crTCA cycle enzymes in tobacco leaf cells, the cell extracts were prepared by adding the enzyme assay buffer to the ground tissue. 1% (v/v) Triton x-100 was also added to aid in chloroplast lysis. Polyvinyl polypyrrolidone (PVPP) was added at 5% (w/v) to remove phenolic compounds. The lysate was centrifuged at 10,000 x g for 20 min, and supernatants were carefully collected. For preparation of cell extracts for HyTh-KOR, all steps and reagents were treated under anaerobic conditions. For the BrBt-SCS samples, the cell extracts were applied to a Ni-NTA spin column (Qiagen) to concentrate the BrBt-SCS proteins and then dialyzed with the same buffer used in the SCS assay. Protein concentration was estimated using the Bradford reagent (BioRad). Enzyme activities were measured spectrophotometrically as described above. In all cases, tobacco transformed with pCAMBIA-EGFP alone was a control for assays and Western blots.

### Generation and growth of transgenic Camelina sativa

The selected crTCA cycle genes were synthesized by GenScript with codon optimization for *C. sativa* onto a pUC57 backbone and flanked by attL/R sites for downstream Gateway cloning into the PC-GW vector series [68]. The genes that constitute the crTCA cycle are cloned in three separate, multi-gene vectors. **Supplementary Table S4** lists each multi-gene vector, its constituent crTCA cycle genes, the att sites used in Gateway cloning (ThermoFisher Scientific), the constitutive promoter, chloroplast transit peptide and terminator. These vectors were assembled using Gateway cloning technology and were positively verified by restriction analysis and sequencing (Eurofins Genomics). crTCA plasmids and empty vector controls were transformed into *Agrobacterium tumefaciens* GV3101 via electroporation. *Camelina sativa* (“Calena”) was transformed via vacuum-assisted floral dip [71].

Seeds collected from the transformed plants (T0) were plated onto 0.5x MS + 1% Sucrose + 0.8% agar plates (Constructs 1 and 2) or 0.5x MS + 1% Sucrose + 0.8% agar + 25 ug/ml BASTA (phosphinothricin) plates (Construct 3). Wild-type plants used in this study are a combined group of WT and null segregants (i.e. siblings of T1 plants that are WT) verified by PCR. Genomic DNA was extracted from ∼50mg of leaf tissue using a CTAB DNA extraction protocol [72] and used for genotyping.

*C. sativa* seeds were stratified on 0.5x MS-agar medium for 3 days and selected for marker expression after 7-9 days of growth in a controlled environment (Percival Scientific). Individual seedlings were transferred to 6” diameter pots in a 1:1 mixture of SunGro Sunshine Mix #8 (SunGro Horticulture) and sterile sand. The substrate mixture was supplemented with 14-14-14 Osmocote fertilizer following manufacturer’s instructions. Positions of the plants were randomized and changed daily. Greenhouse experiments were conducted in March and April with supplemental light providing for a minimum of a 14-hour light period and temperature control for a maximum of 25 °C during the day. The plants were grown in controlled environments: either a Caron 6340 Growth Chamber or in a “C-type” chamber at the NCSU Phytotron. Plant positions were randomized and changed daily and the plants were kept well-watered. The light intensity for the low-light experiment was 200 µmol m^-2^ s^-1^ and 1200 µmol m^-2^ s^-1^ for the high-light experiment. Light period remained constant at 14-hours as well as a day/night temperature of 25/18° C.

### Gas exchange, chlorophyll fluorescence and modelling photosynthesis

Photosynthetic gas exchange and chlorophyll fluorescence were measured using a LI-6400XT portable photosynthesis system (LI-COR Biosciences) equipped with a leaf chamber fluorometer (LI-6400-40). The diurnal changes in photosynthesis were measured on 4-5 week-old, fully-expanded leaves every two hours between 4:00 and 16:00. Light levels inside of the leaf chamber were set to mimic the ambient light intensities throughout the course of the day. Light levels ranged from 0 µmol m^-2^ s^-1^ to 750 µmol m^-2^ s^-1^ with constant 10% blue light.

Relative humidity within the leaf chamber was kept ≥65%. A/Ci response was measured by maintaining saturating light levels (1,800 µmol m^-2^ s^-1^) while changing the concentration of reference CO_2_ and measuring rates of net photosynthesis (Anet). This response was measured first at ambient CO_2_ concentrations before being decreased incrementally to the lowest concentration. The plant was then measured twice at ambient CO_2_ before being increased stepwise to the highest concentration. At all measured concentrations, leaf temperature (25 °C) and relative humidity (65 ± 5%) were kept constant. The A/Ci response was measured in at least 3 individuals per line. Chlorophyll fluorescence measurements were performed simultaneously with gas exchange measurements. Dark-adapted measurements were done at 4am, before dawn. Light-adapted measurements were performed with stable PPFD of 750 µmol m^-2^ s^-1.^ Calculations of chlorophyll fluorescence-derived parameters were done as previously described [73]. All photosynthetic and plant physiology measurements were conducted in at least triplicate for each transgenic or control line. Results presented in tables are mean + standard error of the mean (SEM) for each treatment. A/Ci response was analyzed using the R package *plantecophys* [74] using the “default” fit method for the Farquhar-von Caemmerer-Berry model of C3 photosynthesis and the measured R_d_. Where indicated, statistical differences between treatments were evaluated using ANOVAs and the Tukey HSD method using the R package *agricolae* [75] with a significance threshold of 0.05.

### RNA-seq, differential gene expression and correlation network analysis

Leaf discs of the youngest, fully-expanded leaves of 5-6 week old plants were collected under liquid nitrogen and stored at -80 C until extraction. Total RNA was extracted from pulverized leaf tissue using the PureLink RNA mini kit (ThermoFisher Scientific) following manufacturer’s instructions. Contaminating gDNA was digested out of the samples using the PureLink DNase (ThermoFisher Scientific). Concentrations and purity were determined using both a Qubit 2 fluorometer and a Nanodrop spectrophotometer (ThermoFisher Scientific).

Purified RNA was sent to BGI (https://www.bgi.com/) for cDNA synthesis, library preparation and sequencing. Trimmomatic [76] was used to trim off sequencing adapters and remove low quality reads. Paired-end reads were aligned to the most recent *C. sativa* reference genome using HISAT2 [77]. FeatureCounts [78] was then used to quantify read count per transcript. DESeq2 [79] was then used to analyze differential gene expression between different transgenic lines. Genes were considered to be differentially expressed between WT and transgenic plants if the Bejamini-Hochberg adjusted p-value was ≤ 0.05. Bedtools [80] was used to quantify reads that mapped to crTCA transgenes. For weighted gene co-expression network analysis (WGCNA), we used the full expression dataset with incorporation of the transgene counts. Low abundance (< 5 reads) and low variance (< 10) reads were filtered to reduce noise in the analysis. Hierarchical clustering was performed on expression data using Euclidian distance of samples (hclust(), method = average). An unsigned weighted gene co-expression network was then created using the R package *wgcna* [28] to create modules that correlate to specific crTCA transgene expression. The transgenes that had strong correlation (r > 0.6 or r < -0.6) were combined into a list with their respective strength of correlation. We then performed GO enrichment analysis for both positively and negatively correlated genes. These lists were then analyzed using g:profiler and Cytoscape [81].

### Metabolite extraction and relative quantification

Steady-state metabolite extraction and quantification were performed as previously reported [82] with slight modifications. Briefly, all samples were extracted in a 1:1 (v/v) methanol (MeOH):chloroform solution containing acid washed glass beads at 4°C for 6 hours, with the solution vortexed hourly. 0.5 mL of ddH2O was added and the upper aqueous phase removed and centrifuged in at 0°C. Samples were then frozen, lyophilized and reconstituted in a 1:1 (v/v) MeOH:ddH2O solution. ^13^C-succinic acid was added as an internal standard prior to the extraction and used to normalize samples.

A Thermo Vanquish UHPLC system was used for chromatographic separation and a Orbitrap ID-X MS equipped with electrospray ionization (ESI) source was used for detection of metabolites (ThermoFisher Scientific). For the determination of amino acids, the mobile phase solvents, A and B contained 5% acetonitrile with 0.1% formic acid in water and water with 0.1% formic acid, respectively. A 1 μL sample was injected on to a Waters BEH Amide column Z (2.1 x 100 mm, 1.7 µm, Waters) column that was held at 40°C, and the following gradient was used: the initial concentration of 8% B was linearly increased to 15% B over 1 minute, and to 40% B over the next 8 minutes. The gradient was then brought to 99% B at 8.1 min and was held for an additional 10 minutes before returning to 8% B over 10.1 min. A 12 minute equilibration was used to return the column to the starting conditions (8% B) prior to the next injection. For the detection of the amino acids, a positive ionization mode (ESI+) was used with a scan range of 70-350 m/z. ESI spray voltage was 3.5 kV with the vaporizer temperature set to 350 °C and capillary temperature set to 325 °C. Default gas settings were used: sheath gas 50 au; aux gas 10 au; sweep gas 1 au. The mass resolving power of MS1 was 30000 full width at half maximum (fwhm) at m/z of 200 with a standard automatic gain control target. MS2 data was collected with resolving power of 15000 fwhm, a HCD collision energy of 25, and the cycle time of 0.6 seconds. For the determination of organic acids and sugars, the mobile phase solvents, A and B contained 95% acetonitrile, 5% H2O with 10 mM ammonium bicarbonate and 5% acetonitrile, 95% H2O with 10 mM ammonium bicarbonate, respectively. A 4 μL sample was injected on to a Waters BEH Amide column Z (2.1 x 100 mm, 1.7 µm, Waters) that was held at 45°C, and the following gradient was used: the initial concentration of 1% B was linearly increased to 70% B over 6 minutes, and to 1% B over the next 0.1 minutes. The gradient was then held for an additional 10 minutes at 1% B. For the detection of the organic acids and sugars, a negative ionization mode (ESI-) was used with a scan range of 100-1000 m/z. ESI spray voltage was 3.5 kV with the vaporizer temperature set to 350 °C and capillary temperature set to 325 °C. Default gas settings were used: sheath gas 50 au; aux gas 10 au; sweep gas 1 au. The mass resolving power of MS1 was 60000 fwhm at m/z of 200 with a standard automatic gain control target.

MS2 data was collected with resolving power of 15000 fwhm, a stepped HCD collision energy of 15, 30 and 45, and acquired up to 5 independent scans. For both amino acids and organic acid/sugar methods, raw data files were analyzed in Skyline [83] for peak integration and quantification. Relative quantification was estimated using an external standard calibration curve for all of the amino acids, 2-oxoglutarate, sucrose, fructose and glucose. Succinate, malate, pyruvate and phosphoenolpyruvate were estimated using only one external standard and are therefore shown as relative peak areas rather than a molarity per milligram of fresh weight.

### Metabolic flux analysis

Plant growth and gas exchange methods were used as described previously [84]. The youngest, fully-expanded leaves were used for gas exchange and labeling experiments. A LI-COR 6800 portable photosynthesis system (LI-COR Biosciences) was used to measure carbon assimilation. The reference [CO_2_] was set to 600 ppm, light intensity was 1500 μmol m^−2^ s^−1^, temperature was 22°C, and relative humidity was 70%. After 10-15 min acclimation, the CO_2_ source was switched to ^13^CO_2_ with all other parameters held constant. Gases were mixed with mass flow controllers (Alicat Scientific) controlled by a custom-programmed Raspberry Pi touchscreen monitor (Raspberry Pi foundation; code available upon request). Labeled leaf samples were collected at time points of 0, 0.5, 1, 2, 2.5, 3, 5, 7, 10, 15, 30, and 60 min. Liquid nitrogen was directly sprayed on the leaf surface via a customized fast quenching (0.1-0.5 s to < 0°C) labeling system [84]. The frozen leaf sample was stored at -80°C. Three biological replicates for each time points were collected.

Most C3 cycle intermediates were analyzed by a reverse phase LC-MS/MS method by an ACQUITY UPLC pump system (Waters) coupled with Waters XEVO TQ-S UPLC/MS/MS (Waters). Metabolites were separated by a 2.1×50 mm ACQUITY UPLC BEH C18 Column (Waters) at 40°C. A multi-step gradient was applied with mobile phase A (10 mM tributylamine in 5% (v/v) methanol) and mobile phase B (methanol): 0-1 min, 95-85% A; 1-6 min, 65-40% A; 6-7 min, 40-0% A; 7-8 min, 0% A; 8-9 min, 100% A, at a flow rate of 0.3mL min−1. Mass spectra were acquired using multiple reaction monitoring (MRM) in negative electrospray ionization (ESI) mode as described previously [85] with slight modifications. The source temperature was 120°C and the desolvation temperature was 350°C. Nitrogen was used as a sheath and auxiliary gas and collision gas (argon) was set to 1.1 mTorr. Gas flow for the desolvation and cone was set to 800 and 50 L/h, respectively. The scan time was 0.1 ms.

Other phosphorylated metabolites (e.g., sugar phosphate, 2PG, and PEP) and nucleotide sugars (ADPG and UDPG) were analyzed using an anion-exchange LC-MS/MS method described previously [86] with slight modifications. Metabolites were reconstituted in 100 μL of water from lyophilized extract, and 10 μL of extracts were injected into an ACQUITY UPLC pump system (Waters) coupled with a Xevo ACQUITY TQ Triple Quadrupole Detector (Waters). Metabolites were separated by an IonPac AS11 analytical column (2 × 250 mm, Dionex) equipped with an IonPac guard column AG11 (2 × 50 mm, Dionex) at a flow rate of 0.35 mL min−1. A multi-step gradient was applied with mobile phase A (0.5 mM KOH) and mobile phase B (75 mM KOH): 0-2 min, 100% A; 2-4 min, 100-93% A; 4-13 min, 93-60% A; 13-15 min, 0% A; 15-17 min, 100% A. The KOH concentration was suppressed by a post-column anion self-regenerating suppressor (Dionex ADRS 600, ThermoFisher Scientific), with the current of 50 mA and flow rate of 3.5 mL min−1. An IonPac ATC-3 Anion Trap Column (4 × 35 mm), conditioned with 2M KOH, was used to remove contaminant ions from KOH solvents. Mass spectra were acquired using MRM in negative electrospray ionization (ESI) mode.

Amino acids, organic acids, and sucrose were analyzed using a GC-EI-MS system. Amino acids and organic acids were derivatized by methoximation, followed by tert-butyldimethylsilylation. Sucrose was derivatized by methoximation, followed by trimethylsilylation. Samples were analyzed by an Agilent 7890 GC system (Agilent) coupled to an Agilent 5975C inert XL Mass Selective Detector (Agilent) with an autosampler (CTC PAL) (Agilent). Metabolites were separated by an Agilent VF5ms GC column, 30 m x 0.25 mm x 0.25 m with 10 m guard column (Part number: CP9013, Agilent). For amino acids and organic acids, 1 μL of the derivatized sample was injected in 10 split mode with helium carrier gas with flow rate of 1.2 mL min−1. The oven temperature gradient was: 100 °C (4 min hold), increased at 5°C/min to 200°C, then a 10 °C/min to 320°C, held at 320 ° for 10 min. Electron ionization (EI) is at 70 eV and the mass scan range was 100-600 amu. The ionization source temperature was set at 150 °C and the transfer line temperature 300°C.

Glucose and Fructose were analyzed by GC-CI-MS by an Agilent 7890B GC system (Agilent) coupled to an Agilent 7010B triple quadrupole GC/MS with an autosampler (CTC PAL) (Agilent). An Agilent VF5ms GC column, 30 m x 0.25 mm x 0.25 m with 10 m guard column (Part number: CP9013) was used. One μL of the derivatized sample was injected with helium carrier gas with flow rate of 1.2 mL min^−1^. The oven temperature gradient was: 40 °C (1 min hold), increased at 40 °C/min to 150°C, then a 10 °C/min to 250°C, then a 40 °C/min to 320°C, and finally held at 320 °C for 4.5 min. Chemical ionization (CI) was used and the mass scan range was 150-650 amu with step size 0.1 amu. The ionization source temperature was set at 300 °C and the transfer line temperature was 300°C.

Data from LC-MS/MS were acquired with MassLynx 4.0 (Agilent). Data from GC-EI-MS was acquired with Agilent GC/MSD Chemstation (Agilent). Data from GC-CI-MS was acquired with Agilent MassHunter Workstation (Agilent). Metabolites were identified by retention time and mass to charge ratio (m/z), in comparison with authentic standards. Both LC-MS and GC-MS data were converted to MassLynx format and processed with QuanLynx software for peak detection and quantification. Natural abundance were corrected by Isotopomer Network Compartmental Analysis software package [87] (INCA1.8, http://mfa.vueinnovations.com) implemented in MATLAB 2018b. Mass isotopomer distribution (MID) for each metabolite in which n 13C atoms are incorporated is calculated by the equation of:

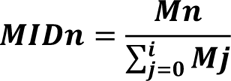

Where *Mn* represents the isotopomer abundance for each metabolite. The ^13^C enrichment of the metabolite possessing *i C* atoms is calculated by the equation of:

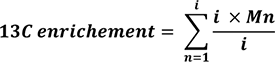

### Protein extraction and peptide preparation

Five biological replicates were included for each line. Leaf tissue from photosynthetically-active leaves was harvested and immediately frozen in liquid nitrogen. This tissue was stored at -80°C until it was ground to a fine powder under liquid nitrogen. The tissue was then weighed into 200 mg aliquots. Alternatively, chloroplasts were isolated from fresh leaf tissue using a commercially available chloroplast isolation kit (Sigma Product #: CPISO) following manufacturer’s instructions. Yield varied between a limited (n = 3) sample set. 200-300 mg of each sample was used for protein extraction in 1 ml of SDT lysis buffer [4% (w/v) SDS, 100 mM Tris-HCl pH 7.6, 0.1 M DTT]. Ground leaf tissue was lysed by bead-beating in lysing matrix E tubes (MP Biomedicals) with a Bead Ruptor Elite (Omni International) for 5 cycles of 45 sec at 6.45 m/s with 1 min dwell time between cycles; followed by heating to 95°C for 10 min. The lysates were centrifuged for 5 min at 21,000 x g to remove cell debris.

Supernatant was used for purification and digestion using the filter-aided sample preparation (FASP) protocol described previously [88]. All centrifugations mentioned below were performed at 14,000 x g. Samples were loaded onto 10 kDa MWCO 500 μl centrifugal filters (VWR International) by combining 60 μl of lysate with 400 μl of Urea solution (8 M urea in 0.1 M Tris/HCl pH 8.5) and centrifuging for 20 min. This step was repeated once to load a total of 120 ul of lysate. Filters were washed once by applying 200 μl of urea solution followed by 20 min of centrifugation to remove any remaining SDS. 100 μl IAA solution (0.05 M iodoacetamide in Urea solution) was then added to filters for a 20 min incubation at room temperature followed by centrifugation for 20 min. The filters were washed three times with 100 uL of urea solution and 20 min centrifugations, followed by a buffer exchange to ABC (50 mM Ammonium Bicarbonate). Buffer exchange was accomplished by three cycles of adding 100 μl of ABC and centrifuging for 20 min. Tryptic digestion was performed by adding 1 μg of MS grade trypsin (ThermoFisher Scientific) in 40 μl of ABC to each filter and incubating for 16 hours in a wet chamber at 37°C. Tryptic peptides were eluted by adding 50 μl of 0.5 M NaCl and centrifuging for 20 min. Peptide concentrations were determined with the Pierce Micro BCA assay (ThermoFisher Scientific) following the manufacturer’s instructions.

### LC-MS/MS

All proteomic samples were analyzed by 1D-LC-MS/MS as described previously [89]. The samples were blocked by treatment in the run sequence to avoid carry over of peptides from the transgenic proteins into the empty vector control samples. Empty vector control samples were run first. 1.4 μg peptide of each sample were loaded with an UltiMate^TM^ 3000 RSLCnano Liquid Chromatograph (ThermoFisher Scientific) in loading solvent A (5 % acetonitrile, 0.05 % trifluoroacetic acid) onto a 5 mm, 300 µm ID C18 Acclaim® PepMap100 pre-column (Thermo Fisher Scientific). Elution and separation of peptides on the analytical column (75 cm x 75 µm analytical EASY-Spray column packed with PepMap RSLC C18, 2 µm material, Thermo Fisher Scientific; heated to 60 °C) was achieved at a flow rate of 300 nl min^-1^ using a 140 min gradient going from 95 % buffer A (0.1 % formic acid) to 31 % buffer B (0.1 % formic acid, 80 % acetonitrile) in 102 min, then to 50 % B in 18 min, to 99 % B in 1 min and ending with 99 % B. The analytical column was connected to a Q Exactive HF hybrid quadrupole-Orbitrap mass spectrometer (ThermoFisher Scientific) via an Easy-Spray source. Eluting peptides were ionized via electrospray ionization (ESI). Carryover was reduced by two wash runs (injection of 20 µl acetonitrile, 99 % eluent B) between sample blocks. MS1 spectra were acquired by performing a full MS scan at a resolution of 60,000 on a 380 to 1600 m/z window. MS2 spectra were acquired using a data dependent approach by selecting for fragmentation the 15 most abundant ions from the precursor MS1 spectra. A normalized collision energy of 25 was applied in the HCD cell to generate the peptide fragments for MS2 spectra. Other settings of the data-dependent acquisition included: a maximum injection time of 100 ms, a dynamic exclusion of 25 sec and exclusion of ions of +1 charge state from fragmentation. About 50,000 MS/MS spectra were acquired per sample.

### Protein identification and quantification

A database containing all protein sequences from *C. sativa* cultivar DH55 (NCBI, RefSeq :GCF_000633955.1), as well as the crTCA vector sequences was used. Sequences of common laboratory contaminants were included by appending the cRAP protein sequence database (http://www.thegpm.org/crap/). The final database contained 48,277 protein sequences and is included in the PRIDE submission (see data access statement) in fasta format. Searches of the MS/MS spectra against this database were performed with the Sequest HT node in Proteome Discoverer version 2.3.0.523 (Thermo Fisher Scientific) as described previously [89]. Peptide false discovery rate (FDR) was calculated using the Percolator node in Proteome Discoverer and only peptides identified at a 5% FDR were retained for protein identification. Proteins were inferred from peptide identifications using the Protein-FDR Validator node in Proteome Discoverer with a target FDR of 5%. For label-free quantification based on peptide area under the curve we used the following nodes and settings in Proteome Discoverer: the “Minora feature” detector node in the processing step. In the consensus step, the .msf files were processed with the “feature mapper” node (maximum allowed retention time shift of 10 min, a mass tolerance of 10 ppm (and a S/N threshold of 5) followed by the “Precursor Ions Quantifier” node. The general quantification setting used Unique + Razor peptides with precursor quantification based on area under the curve.

## Data availability

The mass spectrometry metaproteomics data and protein sequence database have been deposited to the ProteomeXchange Consortium via the PRIDE [90] partner repository. [Reviewers log in at:http://www.ebi.ac.uk/pride; User; Password: vF6TXohj]

## Acknowledgements

We would like to thank Dr. Yu-Chun Liu of the METRIC facility at NCSU for collaboration on steady-state metabolite analysis. The LC-MS/MS measurements of protein samples were made using equipment in the Molecular Education, Technology, and Research Innovation Center (METRIC) at North Carolina State University. This work was supported by the U.S. Department of Energy awards: ARPAe AR-0000207 (C.S-M., C.L., K.L., M.J., X.L., A.G., H.S.) & BER DE-SC0018269 (N.W., Y.X., B.E., S.B., Y.S-H., H.S.). This work was also supported by the U.S. Department of Agriculture National Institute of Food and Agriculture under award No. 2022-67013-36672 (M.K.) and the Novo Nordisk Foundation InRoot project under award No. NNF19SA0059362 (H.S. & M.K.).

## Supporting Information

**Fig S1. Reverse crTCA cycle activity**. Reactions were prepared containing all 5 crTCA cycle enzymes and ferredoxin, but instead of feeding the cycle 2-OG, it was provided with succinate and glyoxylate. The reactions were prepared anaerobically and allowed to proceed for 60 min prior to analysis by LC-MS. Controls consist of a reaction containing all components except ICL, as well as a reaction without succinate

**Fig S2. Comparison of 5-step crTCA cycle and 4-step crTCA cycle**. Reactions were prepared in triplicate at the same time under the same conditions. The only difference was that the 4-step crTCA cycle omitted MaFe OGC. All reaction time points were run on LC-MS. The zero timepoint was subtracted from all samples to remove background. The peak areas of the relevant metabolites were measured using MassHunter software and compared. Statistical significance was evaluated using a two way ANOVA with a p-value <0.05. Only one point was statistically significant, and it is indicated with an asterisk (*). A.) Graphs representing the peak areas for unlabeled succinate (117 m/z), single ^13^C-labeled succinate (118 m/z) double ^13^C-labeled succinate (119 m/z). B.) Graphs representing the peak areas for single ^13^C-labeled 2-OG (146 m/z) and double ^13^C-labeled 2-OG (147 m/z). C.) Graphs representing unlabeled glyoxylate (72 m/z) and single ^13^C-abeled glyoxylate (73 m/z). Due to the low detection of the single labeled glyoxylate, duplicate samples were compared for the 4-step and 5-step cycle comparison instead of triplicate samples.

**Fig S3: Transient tobacco expression western blots of crTCA cycle enzymes**. Tobacco leaf tissue was harvested and ground in liquid nitrogen to produce extracts. Tobacco lysate was purified by nickel IMAC, prior to SDS-PAGE and transfer to PVDF. A.NiHa OSR, NoFa ICL, and BrBT SCS were detected using a penta-his primary antibody. These expression tests were conducted using *N. tabacum*. The empty vector transformed tobacco was used as a negative control, while purified recombinant enzymes were used as positive controls. B. HyTh KOR expression was conducted using *N. benthamiana.* HyTh KOR was detected using an antibody specific for a peptide epitope for the large subunit of the HyTh KOR. Empty vector-transformed tobacco was used as a negative control and purified HyTh KOR was used as a positive control.

**Fig S4: The crTCA cycle transformed into *C. sativa*.** Five enzymes and two co-enzymes that consist of the crTCA cycle. To fully test the cycle’s operation in plant cells, we divided the cycle into multiple components. The core cycle was split (grey line) such that each half would contain a carboxylation reaction. Enzyme abbreviations in red compose Construct 1 (C1), those in green compose Construct 2 (C2), and the co-enzymes in blue compose Construct 3 (C3).

**Fig S5: Expression of the crTCA cycle genes in *Camelina sativa*.** A) Normalized transcript abundances of crTCA transgenes relative to Ubiquitin 3 (UBQ3) as an endogenous reference gene. B) Normalized transcript abundances of RbcS homeologs in *C. sativa*. Transcript abundances were determined using RNA sequencing from leaf tissue collected from 5-6 week old individuals (n = 3 individuals / line). Means are presented ± the standard error of the mean (SEM).

**Fig S6: Plant height over development of crTCA lines grown in the greenhouse.** Plant height was measured over the course of the plant’s development (n = 6 individuals / line). Means are presented ± the standard error of the mean (SEM).

**Fig S7: The A/Ci response of transgenic crTCA lines.** The response of photosynthetic rate (Anet) to increasing intercellular carbon concentration (Ci) (n = 3 plants / line).

**Fig S8: Transient ^13^CO_2_ labelling in all measured metabolites.** Mass isotopomer distribution (MID) of all measured ions are shown as points with error bars (n=3, ± one standard deviation). Nominal masses of M0 mass isotopologues are shown in parentheses.

**Table S1: Gene candidates for the crTCA cycle.**

**Table S2: NiHa OSR and MaFe OGC 4-step cycle activities.** The results are an average of three replicates and error reported is one standard deviation.

**Table S3: *In vitro* LC-MS metabolite quantification under both anaerobic and aerobic conditions.**

**Table S4: Design and elements of crTCA vectors for stable in planta expression.**

**Table S5: Normalized relative protein abundance for crTCA proteins and RbcS.**

**Table S6: GO terms associated with crTCA vector expression determined by weighted gene co-expression network analysis (WGCNA).**

**Table S7: Gas exchange, chlorophyll dynamics and seed yield of C. sativa grown under elevated CO_2_ (1200 ppm) and high light intensity (1200 µmol m^-2^ s^-1^).** Maximum rates of photoassimilation (Anet) and stomatal conductance (g_s_) were analyzed on plants acclimated to a light intensity of 1200 µmol m^-2^ s^-1^ PPFD. Dark respiration (R_d_) and dark-adapted measurements of chlorophyll fluorescence was performed 2 hours before dawn. ΦPSII: quantum yield of PSII; F_v_/F_m_: maximum quantum yield of PSII; NPQ: non-photochemical quenching; Fv’/Fm’: light-adapted quantum yield of PSII. Means are ± the standard error of the mean (SEM), n ≥ 3 plants/ line.

**Table S8: Primers and expression vectors for crTCA cycle gene cloning.**

**File S1: Differentially expressed genes of crTCA lines in *C. sativa*.**

**File S2: Relative abundances of proteins in transgenic *C. sativa* expressing the crTCA cycle.**

**File S3: Parameters for transitions of measured metabolites in multiple reaction monitoring (MRM) with Reverse Phase LC-MS/MS, Anion Exchange LC-MS/MS, selected ion monitoring (SIM) with GC-EI-MS, and GC-CI-MS.**

## References

1. Ray DK, Mueller ND, West PC, Foley JA. Yield Trends Are Insufficient to Double Global Crop Production by 2050. PLoS One. 2013;8: e66428. doi:10.1371/journal.pone.0066428

2. Ort DR, Merchant SS, Alric J, Barkan A, Blankenship RE, Bock R, et al. Redesigning photosynthesis to sustainably meet global food and bioenergy demand. Proceedings of the National Academy of Sciences of the United States of America. National Academy of Sciences; 2015. pp. 8529–8536. doi:10.1073/pnas.1424031112

3. Zhu XG, Long SP, Ort DR. Improving photosynthetic efficiency for greater yield. Annu Rev Plant Biol. 2010;61: 235–261. doi:10.1146/annurev-arplant-042809-112206

4. Weber APM, Bar-Even A. Update: Improving the efficiency of photosynthetic carbon reactions. Plant Physiol. 2019;179: 803–812. doi:10.1104/pp.18.01521

5. Spreitzer RJ, Salvucci ME. R <scp>UBISCO<scp> : Structure, Regulatory Interactions, and Possibilities for a Better Enzyme. Annu Rev Plant Biol. 2002;53: 449–475. Available: https://www.annualreviews.org/doi/abs/10.1146/annurev.arplant.53.100301.135233

6. Erb TJ, Zarzycki J. A short history of RubisCO: the rise and fall (?) of Nature’s predominant CO_2_ fixing enzyme. Current Opinion in Biotechnology. Curr Opin Biotechnol; 2018. pp. 100–107. doi:10.1016/j.copbio.2017.07.017

7. Whitney SM, Birch R, Kelso C, Beck JL, Kapralov M V. Improving recombinant Rubisco biogenesis, plant photosynthesis and growth by coexpressing its ancillary RAF1 chaperone. Proc Natl Acad Sci U S A. 2015;112: 3564–3569. doi:10.1073/pnas.1420536112

8. Parry MAJ, Andralojc PJ, Scales JC, Salvucci ME, Carmo-Silva AE, Alonso H, et al. Rubisco activity and regulation as targets for crop improvement. Journal of Experimental Botany. Oxford Academic; 2013. pp. 717–730. doi:10.1093/jxb/ers336

9. Lin MT, Occhialini A, Andralojc PJ, Parry MAJ, Hanson MR. A faster Rubisco with potential to increase photosynthesis in crops. Nature. 2014;513: 547–550. doi:10.1038/nature13776

10. Batista-Silva W, da Fonseca-Pereira P, Martins AO, Zsögön A, Nunes-Nesi A, Araújo WL. Engineering Improved Photosynthesis in the Era of Synthetic Biology. Plant Communications. Elsevier; 2020. p. 100032. doi:10.1016/j.xplc.2020.100032

11. Kebeish R, Niessen M, Thiruveedhi K, Bari R, Hirsch HJ, Rosenkranz R, et al. Chloroplastic photorespiratory bypass increases photosynthesis and biomass production in Arabidopsis thaliana. Nat Biotechnol. 2007;25: 593–599. doi:10.1038/nbt1299

12. Maier A, Fahnenstich H, von Caemmerer S, Engqvist MK, Weber APM, Flügge UI, et al. Transgenic introduction of a glycolate oxidative cycle into A. Thaliana chloroplasts leads to growth improvement. Front Plant Sci. 2012;3: 38. doi:10.3389/fpls.2012.00038

13. Roell MS, von Borzykowski LS, Westhoff P, Plett A, Paczia N, Claus P, et al. A synthetic C4 shuttle via the β-hydroxyaspartate cycle in C3 plants. Proc Natl Acad Sci U S A. 2021;118: 2021. doi:10.1073/pnas.2022307118

14. Carvalho J de F, Madgwick PJ, Powers SJ, Keys AJ, Lea PJ, Parry MA. An engineered pathway for glyoxylate metabolism in tobacco plants aimed to avoid the release of ammonia in photorespiration. BMC Biotechnol. 2011;11: 1–17. doi:10.1186/1472-6750-11-111

15. South PF, Cavanagh AP, Liu HW, Ort DR. Synthetic glycolate metabolism pathways stimulate crop growth and productivity in the field. Science (80-). 2019;363. doi:10.1126/science.aat9077

16. Dalal J, Lopez H, Vasani NB, Hu Z, Swift JE, Yalamanchili R, et al. A photorespiratory bypass increases plant growth and seed yield in biofuel crop Camelina sativa. Biotechnol Biofuels. 2015;8: 1–22. doi:10.1186/s13068-015-0357-1

17. Shen BR, Wang LM, Lin XL, Yao Z, Xu HW, Zhu CH, et al. Engineering a New Chloroplastic Photorespiratory Bypass to Increase Photosynthetic Efficiency and Productivity in Rice. Mol Plant. 2019;12: 199–214. doi:10.1016/j.molp.2018.11.013

18. Bar-Even A, Noor E, Lewis NE, Milo R. Design and analysis of synthetic carbon fixation pathways. Proc Natl Acad Sci U S A. 2010;107: 8889–8894. doi:10.1073/pnas.0907176107

19. Ducat DC, Silver PA. Improving carbon fixation pathways. Current Opinion in Chemical Biology. Elsevier Current Trends; 2012. pp. 337–344. doi:10.1016/j.cbpa.2012.05.002

20. Schwander T, Von Borzyskowski LS, Burgener S, Cortina NS, Erb TJ. A synthetic pathway for the fixation of carbon dioxide in vitro. Science (80-). 2016;354: 900–904. doi:10.1126/science.aah5237

21. Aoshima M, Igarashi Y. A novel oxalosuccinate-forming enzyme involved in the reductive carboxylation of 2-oxoglutarate in Hydrogenobacter thermophilus TK-6. Mol Microbiol. 2006;62: 748–759. doi:10.1111/j.1365-2958.2006.05399.x

22. Aoshima M. Novel enzyme reactions related to the tricarboxylic acid cycle: Phylogenetic/functional implications and biotechnological applications. Applied Microbiology and Biotechnology. Appl Microbiol Biotechnol; 2007. pp. 249–255. doi:10.1007/s00253-007-0893-0

23. Aoshima M, Igarashi Y. Nondecarboxylating and decarboxylating isocitrate dehydrogenases: Oxalosuccinate reductase as an ancestral form of isocitrate dehydrogenase. J Bacteriol. 2008;190: 2050–2055. doi:10.1128/JB.01799-07

24. Dean AM, Koshland DE. Kinetic Mechanism of Escherichia coli Isocitrate Dehydrogenase. Biochemistry. 1993;32: 9302–9309. doi:10.1021/bi00087a007

25. Kanao T, Kawamura M, Fukui T, Atomi H, Imanaka T. Characterization of isocitrate dehydrogenase from the green sulfur bacterium chlorobium limicola: A carbon dioxide-fixing enzyme in the reductive tricarboxylic acid cycle. Eur J Biochem. 2002;269: 1926– 1931. doi:10.1046/j.1432-1033.2002.02849.x

26. Yamamoto M, Ikeda T, Arai H, Ishii M, Igarashi Y. Carboxylation reaction catalyzed by 2-oxoglutarate:ferredoxin oxidoreductases from Hydrogenobacter thermophilus. Extremophiles. 2010;14: 79–85. doi:10.1007/s00792-009-0289-4

27. Yang Y, Li R, Qi M. In vivo analysis of plant promoters and transcription factors by agroinfiltration of tobacco leaves. Plant J. 2000;22: 543–551. doi:10.1046/j.1365-313X.2000.00760.x

28. Langfelder P, Horvath S. WGCNA: An R package for weighted correlation network analysis. BMC Bioinformatics. 2008;9. doi:10.1186/1471-2105-9-559

29. Wurtzel ET, Vickers CE, Hanson AD, Millar AH, Cooper M, Voss-Fels KP, et al. Revolutionizing agriculture with synthetic biology. Nat Plants. 2019;5: 1207–1210. doi:10.1038/s41477-019-0539-0

30. Eastmond PJ, Graham IA. Re-examining the role of the glyoxylate cycle in oilseeds. Trends in Plant Science. Elsevier; 2001. pp. 72–78. doi:10.1016/S1360-1385(00)01835-5

31. Eisenhut M, Roell MS, Weber APM. Mechanistic understanding of photorespiration paves the way to a new green revolution. New Phytologist. New Phytol; 2019. pp. 1762–1769. doi:10.1111/nph.15872

32. Lehner K, Heldt HW. Dicarboxylate transport across the inner membrane of the chloroplast envelope. BBA - Bioenerg. 1978;501: 531–544. doi:10.1016/0005-2728(78)90119-6

33. Araújo WL, Nunes-Nesi A, Nikoloski Z, Sweetlove LJ, Fernie AR. Metabolic control and regulation of the tricarboxylic acid cycle in photosynthetic and heterotrophic plant tissues. Plant, Cell and Environment. 2012. doi:10.1111/j.1365-3040.2011.02332.x

34. Szecowka M, Heise R, Tohge T, Nunes-Nesi A, Vosloh D, Huege J, et al. Metabolic fluxes in an illuminated Arabidopsis rosette. Plant Cell. 2013;25: 694–714. doi:10.1105/tpc.112.106989

35. Winter H, Robinson DG, Heldt HW. Subcellular volumes and metabolite concentrations in spinach leaves. Planta. 1994;193: 530–535. doi:10.1007/BF02411558

36. Heineke D, Riens B, Grosse H, Hoferichter P, Peter U, Flügge UI, et al. Redox transfer across the inner chloroplast envelope membrane. Plant Physiol. 1991;95: 1131–1137. doi:10.1104/pp.95.4.1131

37. Cedrone F, Ménez A, Quéméneur E. Tailoring new enzyme functions by rational redesign. Current Opinion in Structural Biology. Elsevier Current Trends; 2000. pp. 405–410. doi:10.1016/S0959-440X(00)00106-8

38. De Lorenzo V. Evolutionary tinkering vs. rational engineering in the times of synthetic biology Carmen McLeod, Brigitte Nerlich. Life Sci Soc Policy. 2018;14: 1–16. doi:10.1186/s40504-018-0086-x

39. Dubey A, Verma AK. Enzyme engineering for enzyme activity improvement. Enzymes in Food Biotechnology: Production, Applications, and Future Prospects. Academic Press; 2018. pp. 675–689. doi:10.1016/B978-0-12-813280-7.00039-6

40. Medeiros DB, Martins SCV, Cavalcanti JHF, Daloso DM, Martinoia E, Nunes-Nesi A, et al. Enhanced photosynthesis and growth in atquac1 knockout mutants are due to altered organic acid accumulation and an increase in both stomatal and mesophyll conductance. Plant Physiol. 2016;170: 86–101. doi:10.1104/pp.15.01053

41. Nunes-Nesi A, Araújo W, Fernie A. Targeting mitochondrial metabolism and machinery as a means to enhance photosynthesis. Plant Physiol. 2011;155: 101–107. doi:10.1104/PP.110.163816

42. Timm S, Nunes-Nesi A, Florian A, Eisenhut M, Morgenthal K, Wirtz M, et al. Metabolite profiling in arabidopsis thaliana with moderately impaired photorespiration reveals novel metabolic links and compensatory mechanisms of photorespiration. Metabolites. 2021;11: 391. doi:10.3390/metabo11060391

43. Timm S, Florian A, Fernie AR, Bauwe H. The regulatory interplay between photorespiration and photosynthesis. Journal of Experimental Botany. J Exp Bot; 2016. pp. 2923–2929. doi:10.1093/jxb/erw083

44. Ros R, Muñoz-Bertomeu J, Krueger S. Serine in plants: Biosynthesis, metabolism, and functions. Trends in Plant Science. Elsevier Current Trends; 2014. pp. 564–569. doi:10.1016/j.tplants.2014.06.003

45. Igarashi D, Tsuchida H, Miyao M, Ohsumi C. Glutamate:Glyoxylate aminotransferase modulates amino acid content during photorespiration. Plant Physiol. 2006;142: 901–910. doi:10.1104/pp.106.085514

46. Igarashi D, Miwa T, Seki M, Kobayashi M, Kato T, Tabata S, et al. Identification of photorespiratory glutamate:glyoxylate aminotransferase (GGAT) gene in Arabidopsis. Plant J. 2003;33: 975–987. doi:10.1046/j.1365-313X.2003.01688.x

47. Verslues PE, Kim YS, Zhu JK. Altered ABA, proline and hydrogen peroxide in an Arabidopsis glutamate:glyoxylate aminotransferase mutant. Plant Mol Biol. 2007;64: 205–217. doi:10.1007/s11103-007-9145-z

48. Modde K, Timm S, Florian A, Michl K, Fernie AR, Bauwe H. High serine:glyoxylate aminotransferase activity lowers leaf daytime serine levels, inducing the phosphoserine pathway in Arabidopsis. J Exp Bot. 2017;68: 643–656. doi:10.1093/jxb/erw467

49. Somerville CR, Ogren WL. Photorespiration mutants of Arabidopsis thaliana deficient in serine-glyoxylate aminotransferase activity. Proc Natl Acad Sci. 1980;77: 2684–2687. doi:10.1073/pnas.77.5.2684

50. Busch FA, Sage RF, Farquhar GD. Plants increase CO_2_ uptake by assimilating nitrogen via the photorespiratory pathway. Nat Plants. 2018;4: 46–54. doi:10.1038/s41477-017-0065-x

51. Andrews M, Condron LM, Kemp PD, Topping JF, Lindsey K, Hodge S, et al. Will rising atmospheric CO_2_ concentration inhibit nitrate assimilation in shoots but enhance it in roots of C3 plants? Physiol Plant. 2020;170: 40–45. doi:10.1111/ppl.13096

52. Andrews M, Condron LM, Kemp PD, Topping JF, Lindsey K, Hodge S, et al. Elevated CO_2_ effects on nitrogen assimilation and growth of C3 vascular plants are similar regardless of N-form assimilated. J Exp Bot. 2019;70: 683–690. doi:10.1093/jxb/ery371

53. Bloom AJ, Kasemsap P, Rubio-Asensio JS. Rising atmospheric CO_2_ concentration inhibits nitrate assimilation in shoots but enhances it in roots of C3 plants. Physiol Plant. 2020;168: 963–972. doi:10.1111/ppl.13040

54. Zhao H-L, Chang T-G, Xiao Y, Zhu X-G. Potential Metabolic Mechanisms for Inhibited Chloroplast Nitrogen Assimilation under High CO_2_. Plant Physiol. 2021 [cited 7 Sep 2021]. doi:10.1093/plphys/kiab345

55. Domiciano D, Nery FC, de Carvalho PA, Prudente DO, de Souza LB, Chalfun-Júnior A, et al. Nitrogen sources and CO_2_ concentration synergistically affect the growth and metabolism of tobacco plants. Photosynth Res. 2020;144: 327–339. doi:10.1007/s11120-020-00743-w

56. Igamberdiev AU, Bykova N V., Lea PJ, Gardeström P. The role of photorespiration in redox and energy balance of photosynthetic plant cells: A study with a barley mutant deficient in glycine decarboxylase. Physiol Plant. 2001;111: 427–438. doi:10.1034/j.1399-3054.2001.1110402.x

57. Busch FA. Photorespiration in the context of Rubisco biochemistry, CO_2_ diffusion and metabolism. Plant J. 2020;101: 919–939. doi:10.1111/tpj.14674

58. Aoshima M, Ishii M, Igarashi Y. A novel biotin protein required for reductive carboxylation of 2-oxoglutarate by isocitrate dehydrogenase in Hydrogenobacter thermophilus TK-6. Mol Microbiol. 2004;51: 791–798. doi:10.1046/j.1365-2958.2003.03863.x

59. Fraser ME, James MNG, Bridger WA, Wolodko WT. A detailed structural description of Escherichia coli succinyl-CoA synthetase. J Mol Biol. 1999;285: 1633–1653. doi:10.1006/jmbi.1998.2324

60. Yoon KIS, Ishii M, Igarashi Y, Kodama T. Purification and characterization of 2-oxoglutarate: Ferredoxin oxidoreductase from a thermophilic, obligately chemolithoautotrophic bacterium, Hydrogenobacter thermophilus TK-6. J Bacteriol. 1996;178: 3365–3368. doi:10.1128/jb.178.11.3365-3368.1996

61. Reinscheid DJ, Eikmanns BJ, Sahm H. Characterization of the isocitrate lyase gene from Corynebacterium glutamicum and biochemical analysis of the enzyme. J Bacteriol. 1994;176: 3474–3483. doi:10.1128/jb.176.12.3474-3483.1994

62. Chell RM, Sundaram TK, Wilkinson AE. Isolation and characterization of isocitrate lyase from a thermophilic Bacillus sp. Biochem J. 1978;173: 165–177. doi:10.1042/bj1730165

63. Webb B, Sali A. Comparative protein structure modeling using MODELLER. Curr Protoc Bioinforma. 2016;2016: 5.6.1–5.6.37. doi:10.1002/cpbi.3

64. Bridger WA, Ramaley RF, Boyer PD. [14] Succinyl coenzyme a synthetase from Escherichia coli. [EC 6.2.1.5 Succinate: CoA ligase (SDP)]. Methods Enzymol. 1969;13: 70–75. doi:10.1016/0076-6879(69)13018-9

65. Warren GB, Tipton KF. Pig liver pyruvate carboxylase. Purification, properties and cation specificity. Biochem J. 1974;139: 297–310. doi:10.1042/bj1390297

66. Lu W, Clasquin MF, Melamud E, Amador-Noguez D, Caudy AA, Rabinowitz JD. Metabolomic analysis via reversed-phase ion-pairing liquid chromatography coupled to a stand alone orbitrap mass spectrometer. Anal Chem. 2010;82: 3212–3221. doi:10.1021/ac902837x

67. Pham T V., Murkin AS, Moynihan MM, Harris L, Tyler PC, Shetty N, et al. Mechanism-based inactivator of isocitrate lyases 1 and 2 from Mycobacterium tuberculosis. Proc Natl Acad Sci U S A. 2017;114: 7617–7622. doi:10.1073/pnas.1706134114

68. Dalal J, Yalamanchili R, La Hovary C, Ji M, Rodriguez-Welsh M, Aslett D, et al. A novel gateway-compatible binary vector series (PC-GW) for flexible cloning of multiple genes for genetic transformation of plants. Plasmid. 2015;81: 55–62. doi:10.1016/j.plasmid.2015.06.003

69. Leuzinger K, Dent M, Hurtado J, Stahnke J, Lai H, Zhou X, et al. Efficient agroinfiltration of plants for high-level transient expression of recombinant proteins. J Vis Exp. 2013 [cited 12 Jan 2022]. doi:10.3791/50521

70. Lindbo JA. High-efficiency protein expression in plants from agroinfection-compatible Tobacco mosaic virus expression vectors. BMC Biotechnol. 2007;7: 52. doi:10.1186/1472-6750-7-52

71. Liu X, Brost J, Hutcheon C, Guilfoil R, Wilson AK, Leung S, et al. Transformation of the oilseed crop Camelina sativa by Agrobacterium-mediated floral dip and simple large-scale screening of transformants. Vitr Cell Dev Biol - Plant. 2012;48: 462–468. doi:10.1007/s11627-012-9459-7

72. Clarke JD. Cetyltrimethyl ammonium bromide (CTAB) DNA miniprep for plant DNA isolation. Cold Spring Harb Protoc. 2009;4: pdb.prot5177. doi:10.1101/pdb.prot5177

73. Maxwell1 K, Johnson 2 GN. Chlorophyll fluorescence-a practical guide. J Exp Bot. 2000;51: 659–668.

74. Duursma RA. Plantecophys - An R package for analysing and modelling leaf gas exchange data. PLoS One. 2015;10. doi:10.1371/journal.pone.0143346

75. De Mendiburu F, Simon R. Agricolae - Ten years of an open source statistical tool for experiments in breeding, agriculture and biology. PeerJ. 2015;3. doi:10.7287/peerj.preprints.1404

76. Bolger AM, Lohse M, Usadel B. Trimmomatic: A flexible trimmer for Illumina sequence data. Bioinformatics. 2014;30: 2114–2120. doi:10.1093/bioinformatics/btu170

77. Kim D, Paggi JM, Park C, Bennett C, Salzberg SL. Graph-based genome alignment and genotyping with HISAT2 and HISAT-genotype. Nat Biotechnol. 2019;37: 907–915. doi:10.1038/s41587-019-0201-4

78. Liao Y, Smyth GK, Shi W. FeatureCounts: An efficient general purpose program for assigning sequence reads to genomic features. Bioinformatics. 2014;30: 923–930. doi:10.1093/bioinformatics/btt656

79. Love MI, Huber W, Anders S. Moderated estimation of fold change and dispersion for RNA-seq data with DESeq2. Genome Biol 2014 1512. 2014;15: 1–21. doi:10.1186/S13059-014-0550-8

80. Quinlan AR. BEDTools: The Swiss-Army tool for genome feature analysis. Curr Protoc Bioinforma. 2014;2014: 11.12.1–11.12.34. doi:10.1002/0471250953.bi1112s47

81. Reimand J, Isserlin R, Voisin V, Kucera M, Tannus-Lopes C, Rostamianfar A, et al. Pathway enrichment analysis and visualization of omics data using g:Profiler, GSEA, Cytoscape and EnrichmentMap. Nat Protoc. 2019;14: 482–517. doi:10.1038/s41596-018-0103-9

82. Czajka JJ, Kambhampati S, Tang YJ, Wang Y, Allen DK. Application of Stable Isotope Tracing to Elucidate Metabolic Dynamics During Yarrowia lipolytica α-Ionone Fermentation. iScience. 2020;23. doi:10.1016/j.isci.2020.100854

83. Pino LK, Searle BC, Bollinger JG, Nunn B, MacLean B, MacCoss MJ. The Skyline ecosystem: Informatics for quantitative mass spectrometry proteomics. Mass Spectrometry Reviews. NIH Public Access; 2020. pp. 229–244. doi:10.1002/mas.21540

84. Xu Y, Fu X, Sharkey TD, Shachar-Hill Y, Walker BJ. The metabolic origins of non-photorespiratory CO_2_ release during photosynthesis: A metabolic flux analysis. Plant Physiol. 2021;186: 297–314. doi:10.1093/plphys/kiab076

85. Preiser AL, Fisher N, Banerjee A, Sharkey TD. Plastidic glucose-6-phosphate dehydrogenases are regulated to maintain activity in the light. Biochem J. 2019;476: 1539–1551. doi:10.1042/BCJ20190234

86. Paula Alonso A, Dale VL, Shachar-Hill Y. Understanding fatty acid synthesis in developing maize embryos using metabolic flux analysis. Metab Eng. 2010;12: 488–497. doi:10.1016/j.ymben.2010.04.002

87. Young JD. INCA: A computational platform for isotopically non-stationary metabolic flux analysis. Bioinformatics. 2014;30: 1333–1335. doi:10.1093/bioinformatics/btu015

88. Wiśniewski JR, Zougman A, Nagaraj N, Mann M. Universal sample preparation method for proteome analysis. Nat Methods. 2009;6: 359–362. doi:10.1038/nmeth.1322

89. Mordant A, Kleiner M. Evaluation of Sample Preservation and Storage Methods for Metaproteomics Analysis of Intestinal Microbiomes. Microbiol Spectr. 2021;9. doi:10.1128/spectrum.01877-21

90. Vizcaíno JA, Csordas A, Del-Toro N, Dianes JA, Griss J, Lavidas I, et al. 2016 update of the PRIDE database and its related tools. Nucleic Acids Res. 2016;44: D447–D456. doi:10.1093/nar/gkv1145

