## Supplemental Figures for "Introduction of a condensed, reverse tricarboxylic acid cycle for additional CO_2_ fixation in plants"

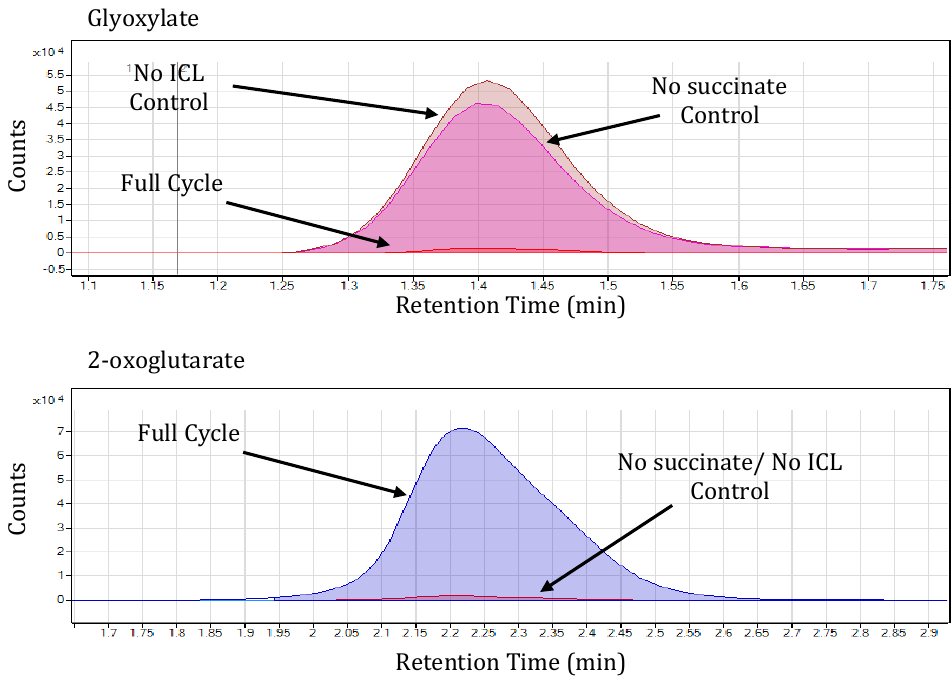
**Figure S1. Reverse crTCA cycle activity**. Reactions were prepared containing all 5 crTCA cycle enzymes and ferredoxin, but instead of feeding the cycle 2-OG, it was provided with succinate and glyoxylate. The reactions were prepared anaerobically and allowed to proceed for 60 min prior to analysis by LC-MS. Controls consist of a reaction containing all components except ICL, as well as a reaction without succinate

**Figure S2. Comparison of 5-step crTCA cycle and 4-step crTCA cycle**. Reactions were prepared in triplicate at the same time under the same conditions. The only difference was that the 4-step crTCA cycle omitted MaFe OGC. All reaction time points were run on LC-MS. The zero timepoint was subtracted from all samples to remove background. The peak areas of the relevant metabolites were measured using MassHunter software and compared. Statistical significance was evaluated using a two way ANOVA with a p-value <0.05. Only one point was statistically significant, and it is indicated with an asterisk (*). A.) Graphs representing the peak areas for unlabeled succinate (117 m/z), single ^13^C-labeled succinate (118 m/z) double ^13^C-labeled succinate (119 m/z). B.) Graphs representing the peak areas for single ^13^C-labeled 2-OG (146 m/z) and double ^13^C-labeled 2-OG (147 m/z). C.) Graphs representing unlabeled glyoxylate (72 m/z) and single ^13^C-abeled glyoxylate (73 m/z). Due to the low detection of the single labeled glyoxylate, duplicate samples were compared for the 4-step and 5-step cycle comparison instead of triplicate samples.

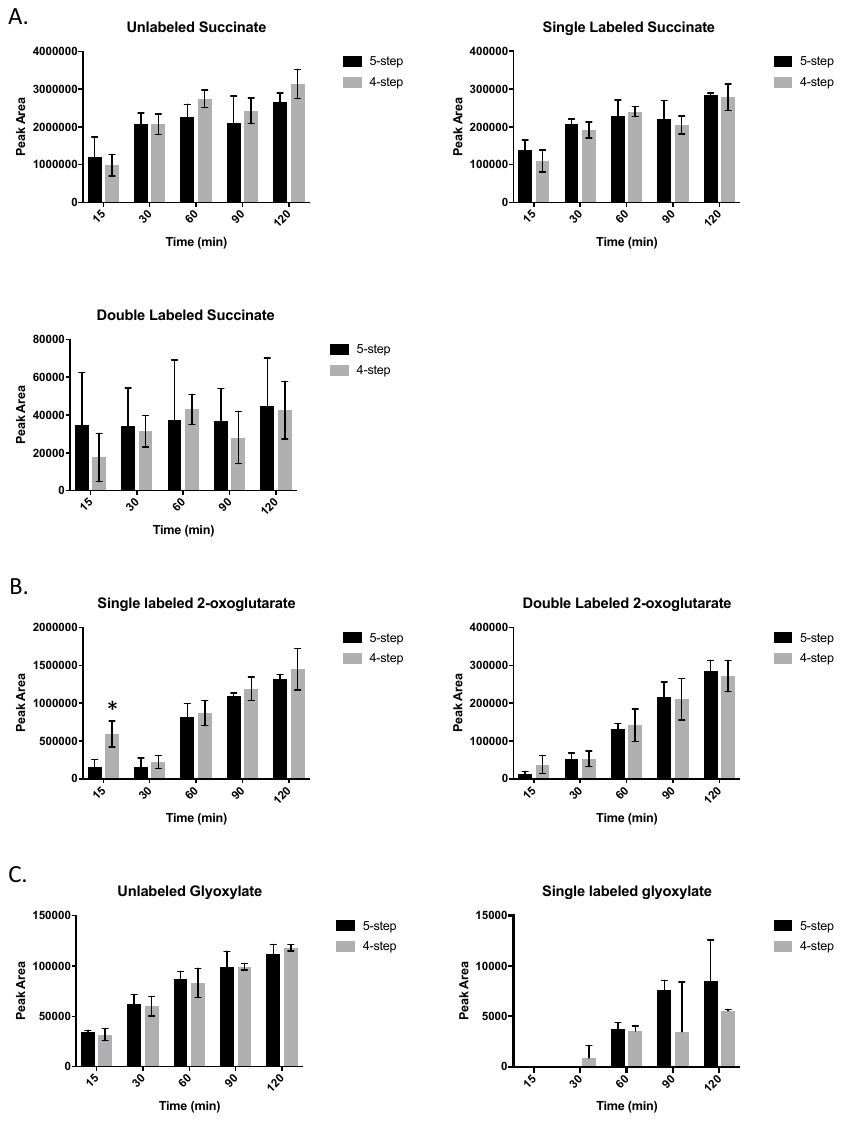

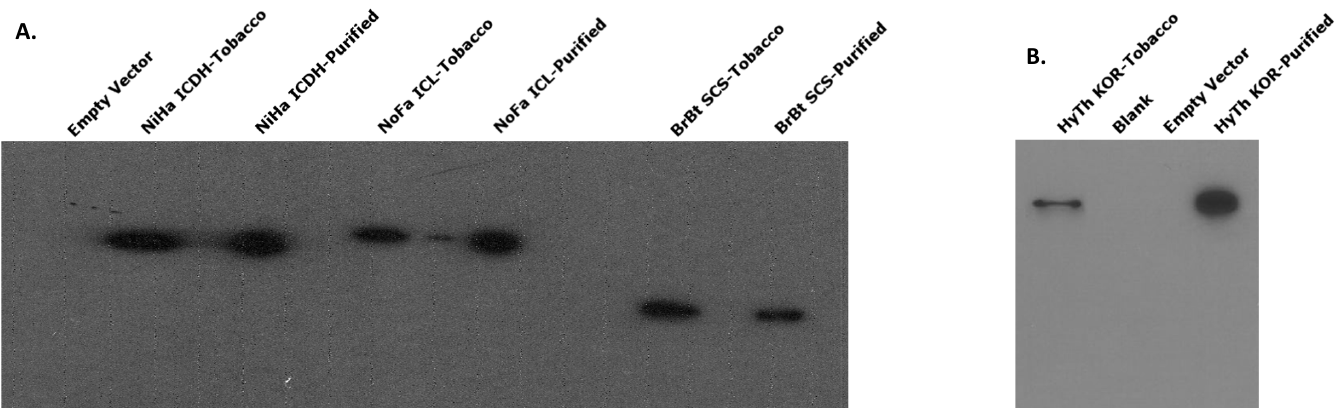
**Figure S3: Transient tobacco expression western blots of crTCA cycle enzymes**. Tobacco leaf tissue was harvested and ground in liquid nitrogen to produce extracts. Tobacco lysate was purified by nickel IMAC, prior to SDS-PAGE and transfer to PVDF. A.NiHa OSR, NoFa ICL, and BrBT SCS were detected using a penta-his primary antibody. These expression tests were conducted using *N. tabacum*. The empty vector transformed tobacco was used as a negative control, while purified recombinant enzymes were used as positive controls. B. HyTh KOR expression was conducted using *N. benthamiana.* HyTh KOR was detected using an antibody specific for a peptide epitope for the large subunit of the HyTh KOR. Empty vector-transformed tobacco was used as a negative control and purified HyTh KOR was used as a positive control.

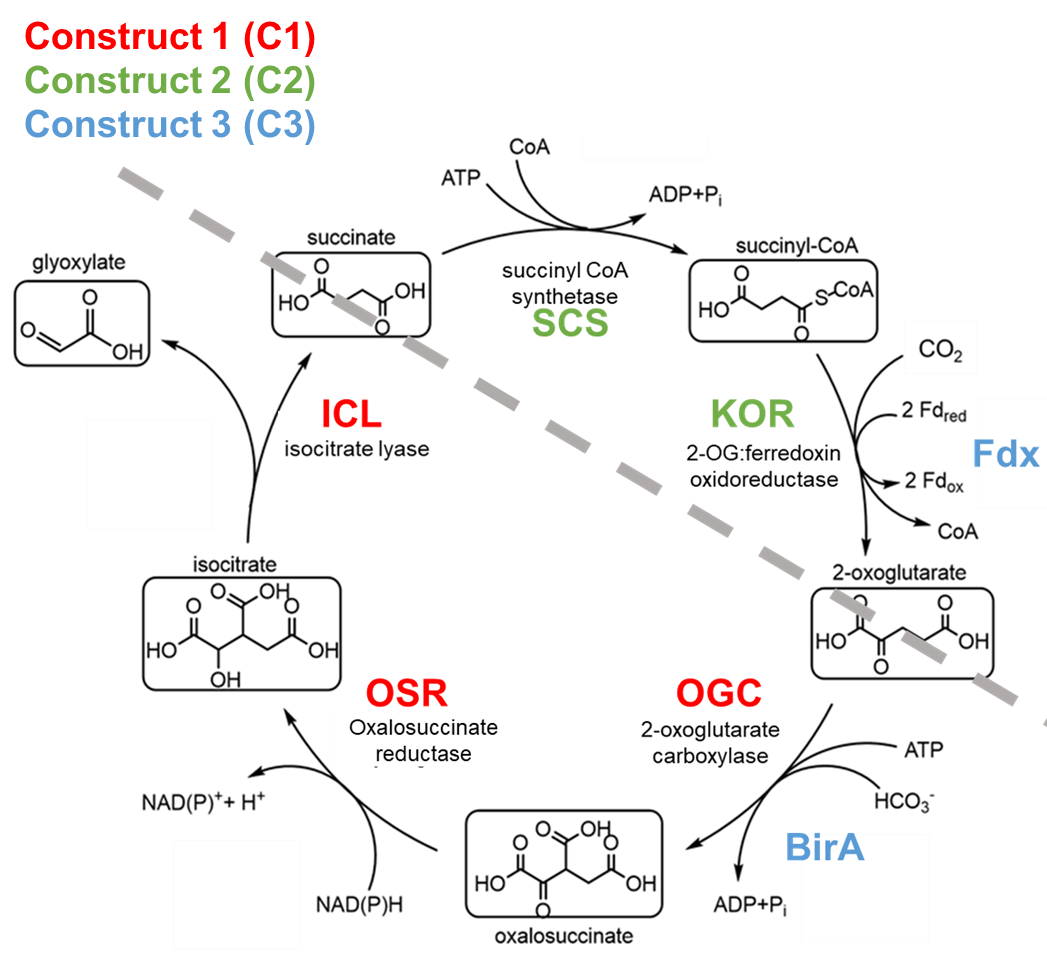
**Figure S4: The crTCA cycle transformed into *C. sativa*.** Five enzymes and two co-enzymes that consist of the crTCA cycle. To fully test the cycle’s operation in plant cells, we divided the cycle into multiple components. The core cycle was split (grey line) such that each half would contain a carboxylation reaction. Enzyme abbreviations in red compose Construct 1 (C1), those in green compose Construct 2 (C2), and the co-enzymes in blue compose Construct 3 (C3).

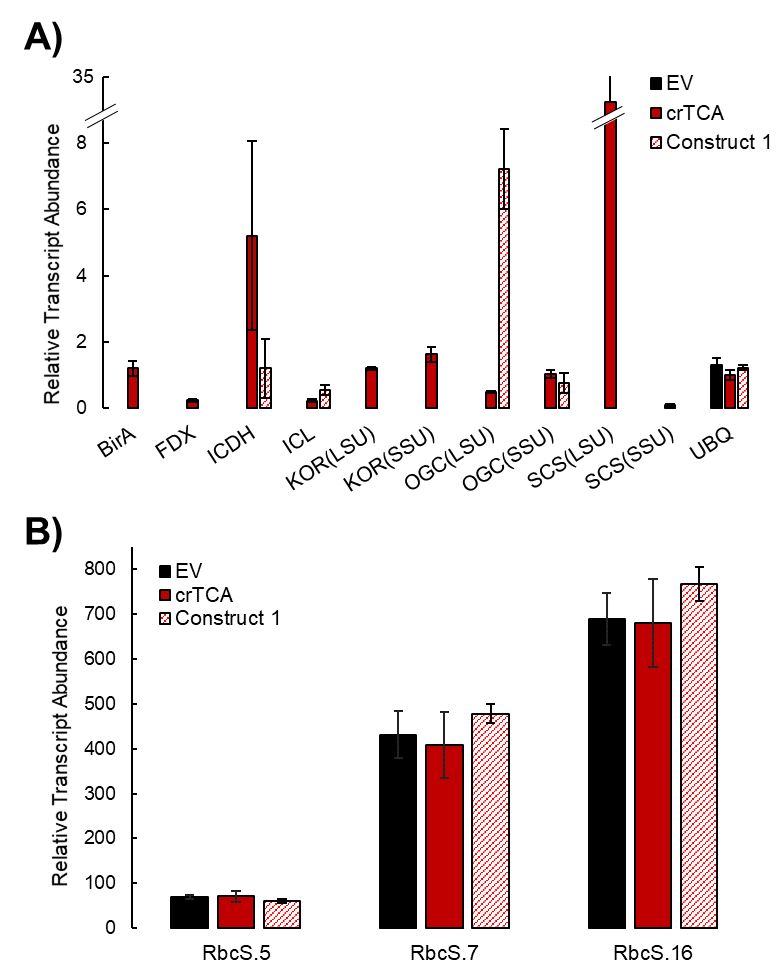
**Figure S5: Expression of the crTCA cycle genes in *Camelina sativa*.** A) Normalized transcript abundances of crTCA transgenes relative to Ubiquitin 3 (UBQ3) as an endogenous reference gene. B) Normalized transcript abundances of RbcS homeologs in *C. sativa*. Transcript abundances were determined using RNA sequencing from leaf tissue collected from 5-6 week old individuals (n = 3 individuals / line). Means are presented ± the standard error of the mean (SEM).

**Figure S6: Plant height over development of crTCA lines grown in the greenhouse.** Plant height was measured over the course of the plant’s development (n = 6 individuals / line). Means are presented ± the standard error of the mean (SEM).

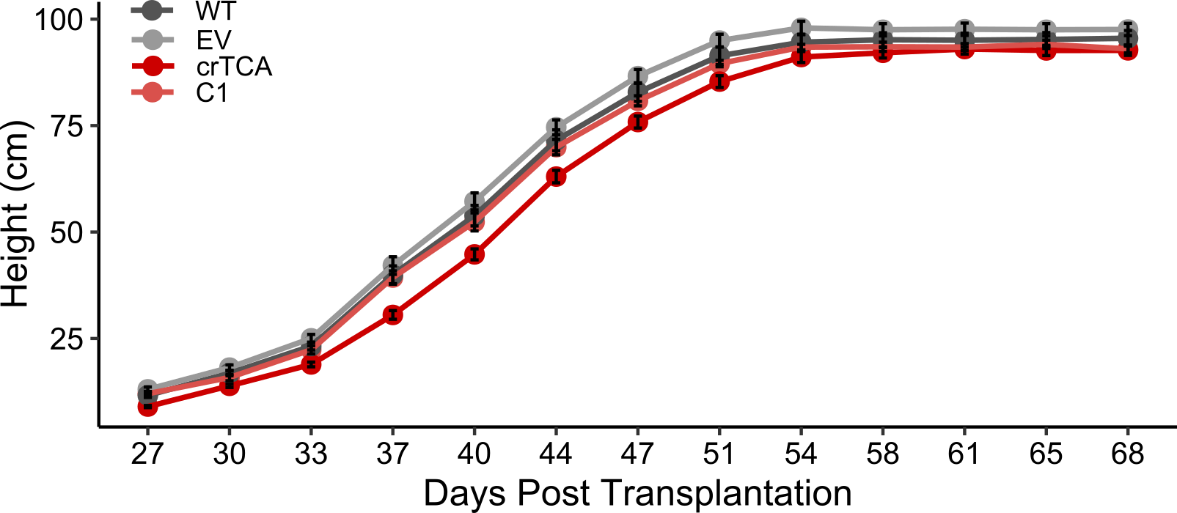

**Figure S7: The A/Ci response of transgenic crTCA lines.** The response of photosynthetic rate (A_net_) to increasing intercellular carbon concentration (C_i_) (n = 3 plants / line).

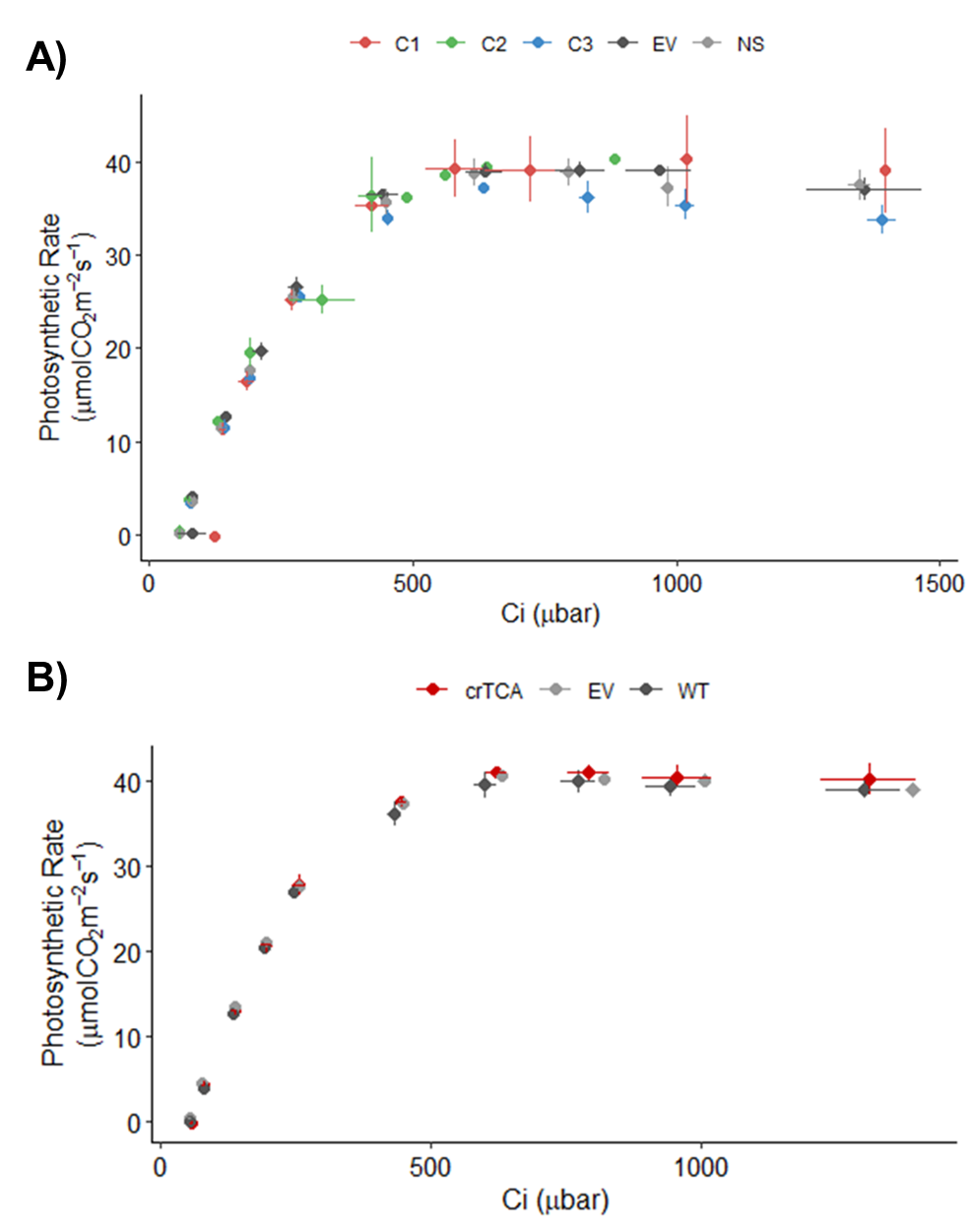

**Figure S8: Transient ^13^CO_2_ labelling in all measured metabolites.** Mass isotopomer distribution (MID) of all measured ions are shown as points with error bars (n=3, ± one standard deviation). Nominal masses of M0 mass isotopologues are shown in parentheses.

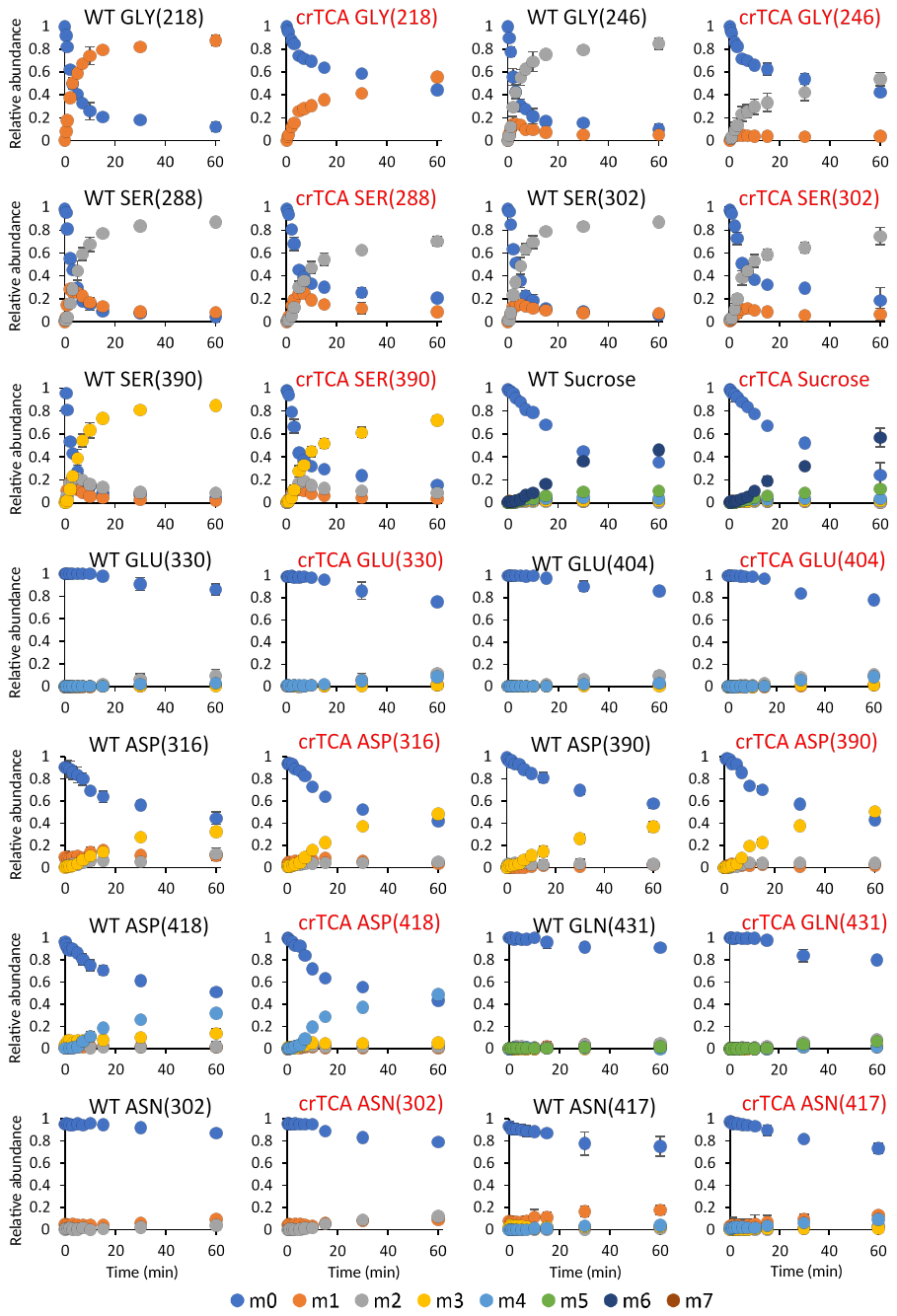

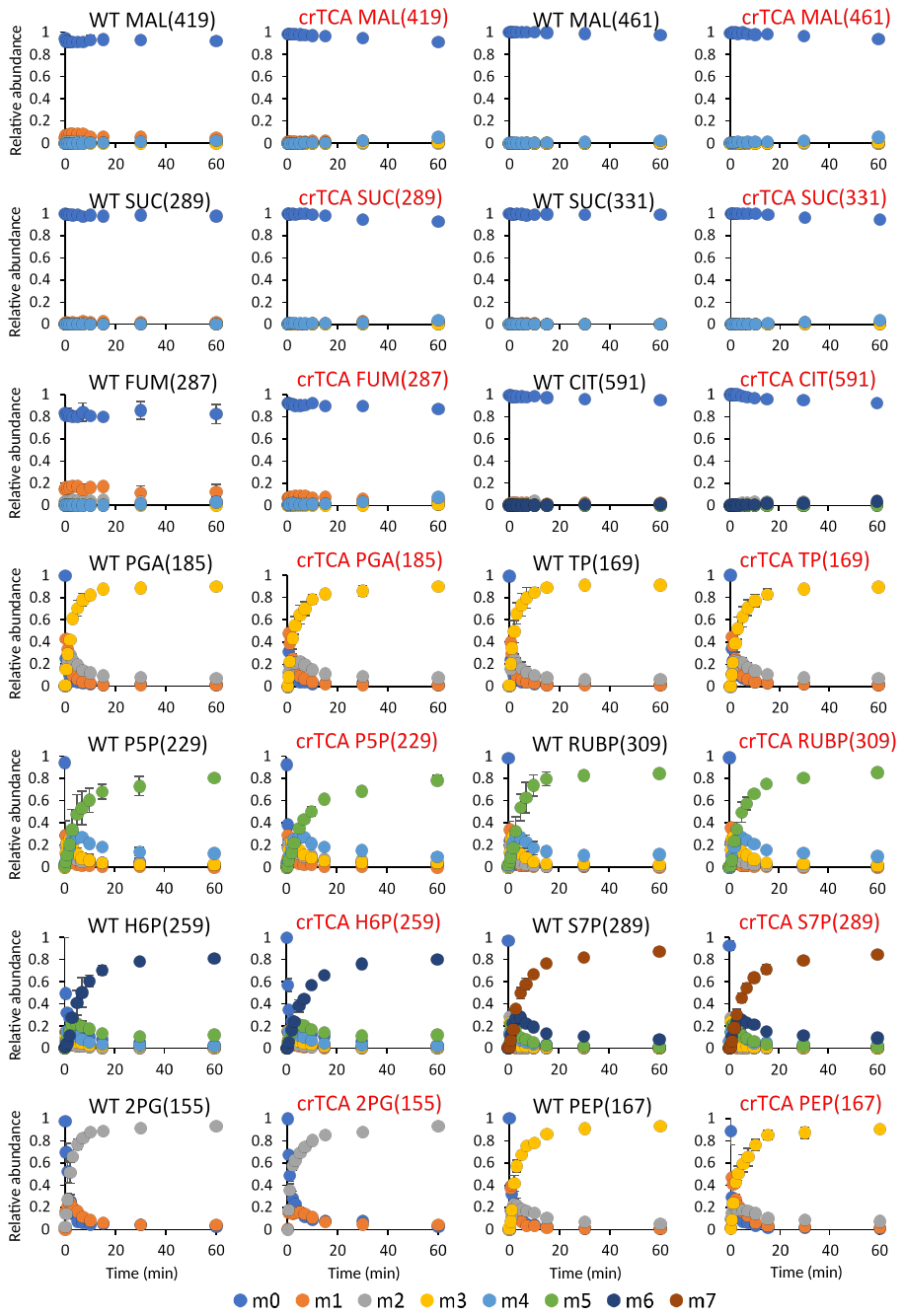

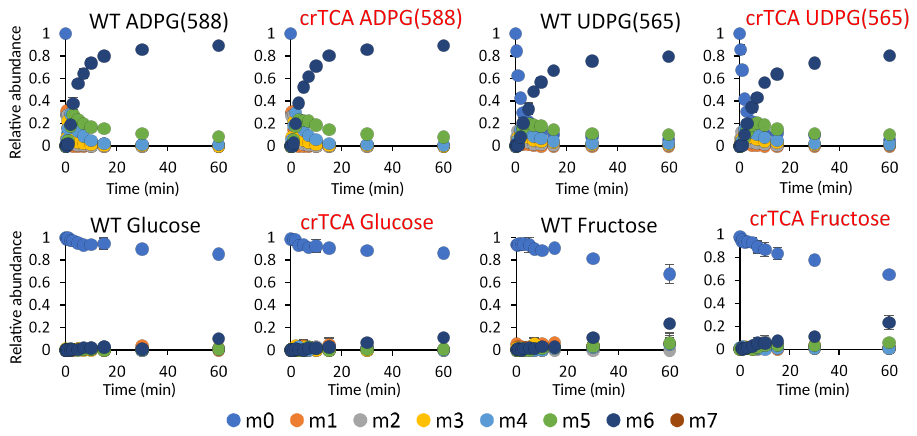
