## Supplemental Tables for "Introduction of a condensed, reverse tricarboxylic acid cycle for additional CO_2_ fixation in plants"

| **Enzyme class** | **Source Organism** | **Protein ID** |
| --- | --- | --- |
| **Step #1**  Succinyl CoA synthetase (SCS) | *Bradyrhizobium* sp. BTAi1 | α: WP_012040734.1 |
|  |  | β: WP_012040733.1 |
|  | *Azotobacter vinelandii* DJ | α: WP_012701526.1 |
|  |  | β: WP_012701527.1 |
|  | *Azospirillum* sp. B510 | α: WP_012975136.1 |
|  |  | β: WP_012975135.1 |
|  | *Escherichia coli* K-12 substr. MG1655 | α: NP_415257.1 |
|  |  | β: NP_415256.1 |
| **Step #2**  2-oxoglutarate: ferredoxin oxidoreductase (KOR) | *Hydrogenobacter thermophilus* TK-6 | α: WP_012963731.1 |
|  |  | β: WP_012963730.1 |
|  | *Bacillus* sp. M3-13 | α: ZP_07708142.1 |
|  |  | β: WP_010193262.1 |
|  | *Haladaptatus paucihalophilus* DX253 | α: WP_007979610.1 |
|  |  | β: WP_007979608.1 |
|  | *Halobacterium* sp. NRC-1 | α: WP_012289282.1 |
|  |  | β: WP_010902808.1 |
|  | *Magnetococcus* sp. MC-1 | α: WP_011713405 |
|  |  | β: WP_011713406.1 |
|  | *Paenibacillus larvae* subsp. larvae B-3650 | α: WP_036654064.1 |
|  |  | β: WP_036654066.1 |
| **Step #3**  2-oxoglutarate carboxylase (OGC) | *Mariprofundus ferrooxydans* PV-1 | α: WP_009849086.1 |
|  |  | β: WP_009849087.1 |
|  | *Hydrogenobacter thermophilus* TK-6 | α: WP_012964024.1 |
|  |  | β: WP_012964023.1 |
|  | *Candidatus Nitrospira defluvii* | α: WP_013247788.1 |
|  |  | β: CBK40961.1 |
|  | *Thiocystis violascens* DSM198 | α: WP_014776649.1 |
|  |  | β: WP_014776651.1 |
|  | *Pseudomonas stutzeri* ATCC14405 | α: ABP77893.1 |
|  |  | β: WP_011911433.1 |
| **Step #4**  Oxalosuccinate reductase (OSR) | *Nitrosococcus halophilus Nc4* | WP_013033479.1 |
|  | *Marine gamma proteobacterium HTCC2080* | WP_007233843.1 |
|  | *Kosmotoga olearia TBF 19.5.1* | WP_015868581.1 |
|  | *Chlorobium limicola* DSMZ245 | BAC00856.1 |
|  | *Acinetobacter baumannii ACICU* | WP_000542119.1 |

**Table S1: Gene candidates for the crTCA cycle**

**Table S2: NiHa OSR and MaFe OGC 4-step cycle activities.** ^a^The results are an average of three replicates and error reported is one standard deviation.

| **Enzyme & assay** | **Specific Activity**  **(U/mg) ^a^** |
| --- | --- |
| NiHa OSR  Carboxylation | 0.24 ± 0.01 |
| MaFe OGC  Pyruvate Carboxylase | 0.11 ± 0.01 |

**Table S3: *In vitro* LC-MS metabolite quantification under both anaerobic and aerobic conditions.**

| **Metabolites** | | **Produced (µM)**  **anaerobic** | **Produced (µM)**  **aerobic** |
| --- | --- | --- | --- |
| glyoxylate | No label | 166 | 148 |
|  | Single ^13^C labeled | 59 | 7.1 |
|  | Double ^13^C labeled | 5.2 | 1.9 |
| succinate | Single ^13^C labeled | 180 | 158 |
|  | Double ^13^C labeled | 42 | 18 |
| oxoglutarate | Single ^13^C labeled | 193 | 65 |
|  | Double ^13^C labeled | 74 | 19 |
|  | Triple ^13^C labeled | 11 | 4.4 |

**Table S4: Design and elements of crTCA vectors for stable *in planta* expression.**

| **Construct** | **Element** | **Enzyme** | **att sites** | **Promoter** | **Transit Peptide** | **Terminator** |
| --- | --- | --- | --- | --- | --- | --- |
| 1 | 1 | OGC SSU | L1, R5 | 35S (in pCAMBIA) | RBCS | 35S |
|  | 2 | OGC LSU | L5, L4 | Act2 | CTP | OCS |
|  | 3 | ICDH | R4, R3 | 35S | RBCS | 35S |
|  | 4 | ICL | L3, L2 | Entcup4 | BCCP | NOS (in pCAMBIA) |
| 2 | 1 | SCS LSU | L1, R5 | 35S (in pCAMBIA) | RBCS | 35S |
|  | 2 | SCS SSU | L5, L4 | Entcup4 | CTP | NOS |
|  | 3 | KOR LSU | R4, R3 | Act2 | BCCP | 35S |
|  | 4 | KOR SSU | L3, L2 | 35S | CTP | NOS (in pCAMBIA) |
| 3 | 1 | BirA | N/A | EntCUP4 | RBCS | OCS |
|  | 2 | Fdx | N/A | Actin2 | RBCS | NOS |

**Table S5: Normalized relative protein abundance for crTCA proteins and RbcS.**

|  | **RbcS**  **(16)** | **RbcS**  **(7)** | **ICDH** | **ICL** | **SCS**  **(SSU)** | **SCS**  **(LSU)** | **OGC**  **(SSU)** |
| --- | --- | --- | --- | --- | --- | --- | --- |
| **Empty Vector** |  |  |  |  |  |  |  |
| Sample 1 | 1.701 | 0.7106 |  |  |  |  |  |
| Sample 2 | 11.37 | 5.301 |  |  |  |  |  |
| Sample 3 | 13.49 | 6.059 |  |  |  |  |  |
| Sample 4 | 11.14 | 4.994 |  |  |  |  |  |
| Sample 5 | 12.30 | 5.025 |  |  |  |  |  |
| Chloroplast | 11.16 | 4.810 |  |  |  |  |  |
| **crTCA** |  |  |  |  |  |  |  |
| Sample 1 | 15.09 | 5.98 | 0.0009 | 0.0068 | 0.8823 | 0.0299 | n.d. |
| Sample 2 | 14.52 | 6.830 | 0.0009 | 0.0106 | 0.0042 | 0.0053 | n.d. |
| Sample 3 | 12.42 | 5.81 | 0.0027 | 0.0207 | 0.0099 | 0.0180 | n.d. |
| Sample 4 | 13.20 | 6.27 | 0.0038 | 0.0234 | 0.0096 | 0.0247 | n.d. |
| Sample 5 | 11.37 | 5.33 | 0.0027 | 0.0219 | 0.0074 | 0.0216 | n.d. |
| Chloroplast | 0.42 | 0.13 | 0.0025 | 0.0014 | 0.0003 | 0.0019 | 0.0004 |
| **Construct 1** |  |  |  |  |  |  |  |
| Sample 1 | 9.118 | 0.8787 | 0.0754 | 0.0033 |  |  | 0.0009 |
| Sample 2 | 11.84 | 5.322 | 0.1067 | 0.0079 |  |  | 0.0127 |
| Sample 3 | 15.19 | 5.523 | 0.1321 | 0.0043 |  |  | 0.0072 |
| Sample 4 | 13.44 | 6.607 | 0.1779 | 0.0076 |  |  | 0.0162 |
| Sample 5 | 13.09 | 5.879 | 0.1575 | 0.0071 |  |  | 0.0153 |
| Chloroplast | 0.0065 | 0.0020 | 0.0004 | 0.0001 |  |  | < 0.0001 |

**Table S6: GO terms associated with crTCA vector expression determined by weighted gene co-expression network analysis (WGCNA).**

| **Construct**  **Association** | **Term Name** | **Term ID** | **padj** | **Pos/Neg** |
| --- | --- | --- | --- | --- |
| **C1** | Organic acid metabolic process | GO:0006082 | 7.812x10^-38^ | Negative |
| **C1** | Carboxylic acid metabolic process | GO:0019752 | 1.808x10^-36^ | Negative |
| **C1** | Oxoacid metabolic process | GO:0043436 | 5.248x10^-35^ | Negative |
| **C1** | Small molecule metabolic process | GO:0044281 | 1.261x10^-33^ | Negative |
| **C1** | Response to cadmium ion | GO:0046686 | 6.526x10^-25^ | Negative |
| **C1** | Response to inorganic substance | GO:0010035 | 6.003x10^-25^ | Negative |
| **C1** | Lipid homeostasis | GO:0055088 | 4.929x10^-2^ | Positive |
| **C1** | Cellular modified amino acid biosynthetic process | GO:0042398 | 3.042x10^-2^ | Positive |
| **C1** | Regulation of circadian rhythm | GO:0042752 | 3.807x10^-2^ | Positive |
| **C1** | Anther wall tepetum development | GO:0048658 | 1.654x10^-2^ | Positive |
| **C2** | Photosynthesis | GO:0015979 | 2.452x10^-55^ | Negative |
| **C2** | Generation of precursor metabolites and energy | GO:0006091 | 3.367x10^-32^ | Negative |
| **C2** | Small molecule metabolic process | GO:0044281 | 2.714x10^-23^ | Negative |
| **C2** | Photosynthesis, light harvesting | GO:0009765 | 1.035x10^-22^ | Negative |
| **C2** | Photosynthesis, light reaction | GO:0019684 | 3.599x10^-39^ | Negative |
| **C2** | Carboxylic acid metabolic process | GO:0019752 | 3.174x10^-21^ | Negative |

**Table S6 (continued)**

| **C2** | Monocarboxylic acid metabolic process | GO:0032787 | 2.138x10^-19^ | Negative |
| --- | --- | --- | --- | --- |
| **C2** | Oxoacid metabolic process | GO:0043436 | 3.905x10^-20^ | Negative |
| **C2** | Carboxylic acid biosynthetic process | GO:0046394 | 1.142x10^-17^ | Negative |
| **C2** | Fatty acid metabolic process | GO:0006631 | 1.235x10^-16^ | Negative |
| **C2** | Cellular catabolic process | GO:0044248 | 1.692x10^-16^ | Positive |
| **C2** | Organic substance catabolic process | GO:1901575 | 1.310x10^-15^ | Positive |
| **C2** | Catabolic process | GO:0009056 | 5.381x10^-15^ | Positive |
| **C2** | Small molecular catabolic process | GO:0044282 | 1.966x10^-14^ | Positive |
| **C3** | Oxidation-reduction process | GO:0055114 | 7.937x10^-14^ | Negative |
| **C3** | Detoxification of nitrogen compound | GO:0051410 | 2.842x10^-1^ | Positive |

**Table S7: Gas exchange, chlorophyll dynamics and seed yield of *C. sativa* grown under elevated CO2 (1200 ppm) and high light intensity (1200 µmol m^-2^ s^-1^) .** Maximum rates of photoassimilation (A_net_) and stomatal conductance (g_s_) were analyzed on plants acclimated to a light intensity of 1200 µmol m^-2^ s^-1^ PPFD. Dark respiration (R_d_) and dark-adapted measurements of chlorophyll fluorescence was performed 2 hours before dawn. ΦPSII: quantum yield of PSII; F_v_/F_m_: maximum quantum yield of PSII; NPQ: non-photochemical quenching; F_v’_/F_m’_: light-adapted quantum yield of PSII. Means are ± the standard error of the mean (SEM) , n ≥ 3 plants / line.

|  | **WT** | **crTCA** | **Construct 1** |
| --- | --- | --- | --- |
| **A_net_ (µmol CO_2_ m^-2^ s^-1^)** | 23.1 ± 0.3^c^ | **24.0 ± 0.1^b^** | **26.7 ± 0.4^a^** |
| **g_s_ (µmol CO_2_ m^-2^ s^-1^)** | 0.31 ± 0.01^c^ | **0.52 ± 0.01^a^** | **0.41 ± 0.01^b^** |
| **R_d_ (µmol CO_2_ m^-2^ s^-1^)** | -2.4 ± 0.01^b^ | **3.2 ± 0.05^a^** | **-2.9 ± 0.1^a^** |
| **ΦPSII** | 0.240 ± 0.02^a^ | 0.219 ± 0.03^a^ | 0.209 ± 0.02^a^ |
| **F_v_/F_m_** | 0.794 ± 0.004^a^ | 0.804 ± 0.015^a^ | 0.811 ± 0.006^a^ |
| **NPQ** | 1.58 ± 0.2^a^ | 1.33 ± 0.4^a^ | 1.81 ± 0.3^a^ |
| **F_v’_/F_m’_** | 0.521 ± 0.01^a^ | 0.552 ± 0.03^a^ | 0.525 ± 0.01^a^ |
| **Total Yield (g / plant)** | 6.18 ± 1.2^a^ | 6.79 ± 0.18^a^ | 6.44 ± 0.1^a^ |
| **Seed Weight (mg / 50 seed)** | 67.1 ± 6.0^a^ | 67.5 ± 3.5^a^ | 64.4 ± 3.2^a^ |

**Table S8: Primers and expression vectors for crTCA cycle gene cloning.**

| Gene name | Organism source | Expression Vector | Forward primer (5’-3’) | Reverse primer (5’-3’) | Tm |
| --- | --- | --- | --- | --- | --- |
| AzB5 (SCS) | *Azospirillum* sp. B510 | pQE1 | ATGAACATCCATGAATACCAG | ACAATTTCACACAGGAAACAGCTA | 65^o^C |
| AzVi (SCS) | *Azotobacter vinelandii* DJ | pQE1 | ATGAATCTGCATGAATACCAGGGC | ACAATTTCACACAGGAAACAGCTA | 65^o^C |
| BrBT (SCS) | *Bradyrhizobium* sp. BTAi1 | pQE1 | ATGAACATCCACGAATACCA | AACGACGGCCAGTGAATTCGAGC | 65^o^C |
| BrBT (SCS) | *Bradyrhizobium* sp. BTAi1 | pET21b | CCGATGACCCCATATGAACATCCACGAATACCA | CGAATGCATCTAGATGACTCGAGTCCGCTCTTCAGTTTTTCAACCAG | 70^o^C |
| EsCo (SCS) | *Escherichia coli* K-12 substr. MG1655 | pQE1 | ATGAACCTGCACGAATACCAAG | AACGACGGCCAGTGAATTCGAGC | 65^o^C |
| BaM3 (KOR) | *Bacillus* sp. M3-13 | pQE1 | ATGATTAACCAACTGTCCTGGAA | ACAATTTCACACAGGAAACAGCTA | 65^o^C |
| BaM3 (KOR) | *Bacillus* sp. M3-13 | pET21b | GCATCTAGATGACCCCATATGATTAAC | ATCCGATGACTCGAGTCCCATAAATTCC | 65^o^C |
| BaM3 (KOR) | *Bacillus* sp. M3-13 | pET28a | GCATCTAGATGACCCCATATGATTAAC | ATCCGATGACTCGAGTTACATAAATTCC | 65^o^C |
| HaNR (KOR) | *Halobacterium* sp. NRC-1 | pQE1 | ATGCCGTACTGGAGCACCGCTGGCC | AACGACGGCCAGTGAATTCGAGC | 65^o^C |
| HaPa (KOR) | *Haladaptatus paucihalophilus* DX253 | pQE1 | ATGCAGGATCTGAACTGGGC | ACAATTTCACACAGGAAACAGCTA | 65^o^C |
| HyTh (KOR) | *Hydrogenobacter thermophilus* TK-6 | pQE1 | ATGGCGTTTGACCTGACGATTAAGATTG | AACGACGGCCAGTGAATTCGAGC | 65^o^C |
| HyTh (KOR) | *Hydrogenobacter thermophilus* TK-6 | pET21b | GGGATCCGATGTCTCATATGGCGTTT | TCGCGAATGACTCGAGATCACAAATTCCCA | 70^o^C |
| HyTh (KOR) | *Hydrogenobacter thermophilus* TK-6 | pET28a | GGGATCCGATGTCTCATATGGCGTTT | TCGCGAATGCCTCGAGATCCCAAATTTACA | 65^o^C |
| MaMC (KOR) | *Magnetococcus* sp. MC-1 | pQE1 | ATGGAAAAGAAGGACCTGA | ACAATTTCACACAGGAAACAGCTA | 65^o^C |
| PaLa (KOR) | *Paenibacillus larvae* subsp. larvae B-3650 | pQE1 | ATGATTAGCCAACTGAGCT | ACAATTTCACACAGGAAACAGCTA | 65^o^C |
| PaLa (KOR) | *Paenibacillus larvae* subsp. larvae B-3650 | pET21b | GCATCTAGATGACCCCATATGATTAGC | GGATCCGATGACTCGAGTGGCTTAAA | 65^o^C |
| PaLa (KOR) | *Paenibacillus larvae* subsp. larvae B-3650 | pET28a | GCATCTAGATGACCCCATATGATTAGC | GGGATCCGATGACTCGAGTTACTTA | 60^o^C |
| NiDe  (OGC) | *Canditatus Nitrospira defluvii* | pQE1 | ATGTTCCGTAAAATCCTGATCG | ACAATTTCACACAGGAAACAGCTATGAC | 67^o^C |
| HyTh  (OGC) | *Hydrogenobacter thermophilus TK-6* | pQE1 | ATGTTCAAAAAAGTCCTGGTCGC | ACAATTTCACACAGGAAACAGCTATGAC | 71.4^o^C |
| ThVi  (OGC) | *Thiocystis violascens DSM198* | pQE1 | ATGCTGCGTAAGATCCTGATTGCGAA | ACAATTTCACACAGGAAACAGCTATGAC | 71.9^o^C |

**Table S8 (continued)**

| MaFe  (OGC) | *Mariprofundus ferrooxydans*  *PV-1* | pQE1 | ATGTTTAAGCGTATCCTGGTGGC | ACAATTTCACACAGGAAACAGCTA | 70^o^C |
| --- | --- | --- | --- | --- | --- |
| PsSt  (OGC) | *Pseudomonas stutzeri ATCC14405* | pQE1 | ATGCGTATCAATGACTTCCGTATCGTC | ACAATTTCACACAGGAAACAGCTATGAC | 71.4^o^C |
| ChLi (OSR) | *Chlorobium limicola* DSM245 | pQE1 | ATGGCCTCCAAGTCCACGATTAT | AACGACGGCCAGTGAATTCGAGC | 66^o^C |
| NiHa (OSR) | *Nitrosococcus halophilus* NC4 | pQE1 | ATGGCCTACGACAAGATTAGCCT | ACGACGGCCAGTGAATTCGAG | 66^o^C |
| KoOl (OSR) | *Kosmotoga olearia TBF 19.5.1* | pQE1 | ATGGAAGGTCAAAAGATTAAAGTG | ACGACGGCCAGTGAATTCGAG | 66^o^C |
| MgPr (OSR) | Marine gamma proteobacterium  HTCC2080 | pQE1 | ATGTCATACAAGCATATTAAAGTCC | ACAATTTCACACAGGAAACAGCTA | 62^o^C |
| CoGl (ICL) | *Corynebacterium glutamicum* ATCC | pQE1 | ATGTCAAATGTCGGCAAACC | ACGACGGCCAGTGAATTCGAG | 64^o^C |
| GoAl (ICL) | *Gordonia alkanivorans* NBRC 16433 | pQE1 | ATGAGCAATGTGGGCAAACC | ACAATTTCACACAGGAAACAGCTA | 62^o^C |
| NoFa (ICL) | *Nocardia farcinica* IFM 10152 | pQE1 | ATGAGCACCACCGGCACCCC | AACGACGGCCAGTGAATTCGAGC | 66^o^C |
| RdPy (ICL) | *Rhodococcus pyridinivorans* AK37 | pQE1 | ATGAGCACGACCGGCACCCC | AACGACGGCCAGTGAATTCGAGC | 66^o^C |
| HyTh-fdx | *Hydrogenobacter thermophilus* TK-6 | pQE1 | ATGGCTCTGCGCACGATG | ACAATTTCACACAGGAAACAGCTA | 58^o^C |
